# Genetic differences in plasticity across environmental scales determine fitness along an ecological gradient

**DOI:** 10.1101/2025.03.19.644087

**Authors:** Greg M. Walter, Giuseppe Emma, Delia Terranova, James Clark, Salvatore Cozzolino, Simon J. Hiscock, Antonia Cristaudo, Jon Bridle

## Abstract

When populations suffer reduced fitness in novel environments, genotypes that better adjust their phenotype to cope with environmental change can aid persistence by reducing the severity of fitness declines. However, we know little about how plastic changes in phenotype allow different genotypes to track environmental variation across ecological gradients, particularly as environments become novel. We transplanted numerous clones of 19 genotypes of a Sicilian daisy, *Senecio chrysanthemifolius*, at four elevations on Mt Etna. We assessed fitness at native and novel elevations and quantified leaf plasticity among and within elevations. Genotypes with higher fitness at novel elevations showed lower variance in fitness, lower plasticity across elevations, but higher plasticity within elevations compared to those with higher fitness in the native range. Our results suggest that there are genotypes hidden in a population whose plasticity better tracks novel environmental variation at multiple scales, which will be crucial for population persistence under rapid environmental change.

## INTRODUCTION

There is an urgent need to understand the capacity for natural populations to produce adaptive responses to ongoing global change (Martin *et al*. 2023; Urban *et al*. 2024). Adaptive phenotypic plasticity, defined as the ability of genotypes to express different beneficial phenotypes as the environment changes, allows populations to maintain fitness as the environment varies (de Jong 1995; Via *et al*. 1995; Sultan 2000; Charmantier *et al*. 2008). However, maintaining fitness via adaptive plasticity becomes difficult when novel environments are encountered, such as when new habitats are colonised, or when environmental change is rapid and unpredictable. This is because adaptive plasticity should only evolve to buffer familiar variation in the environment, meaning that plastic responses shaped by current or historical environments should result in fitness (and population) declines under novel conditions (Bradshaw 1991; Ghalambor *et al*. 2007; Matesanz *et al*. 2010; Reed *et al*. 2010; Fierst 2011; Snell-Rood *et al*. 2018). Population persistence should then be determined by the extent to which genotypes vary in their plastic response to novel environments, and how such differences in plasticity can maintain high enough mean fitness to avoid extinction (Chevin & Bridle 2025).

Within populations, genotypes often vary in plasticity and therefore in their sensitivity to the environment (Bradshaw 1965; Schlichting 1986; Scheiner 1993). Predicting population persistence under global change then requires identifying when and how genetic differences in plasticity can help populations maintain fitness as environments become novel (Yeh & Price 2004; Morris 2014). However, the need to assay hundreds of genotypes means that field studies quantifying genetic variation in plasticity are rare, and limited to a single model species and a few select environments (Matesanz *et al*. 2010; Gianoli & Valladares 2012; Merilä & Hendry 2014; Peschel *et al*. 2020). Consequently, field experiments typically lack the capacity to determine how genetic variation in plasticity determines fitness along environmental gradients. We therefore do not know how genotypes that differ in plasticity make phenotypic adjustments to maintain fitness within native environments, or how the same plasticity determines fitness as environments become novel. This gap in knowledge remains a significant barrier to understanding the adaptive potential of populations facing rapid global change (Chevin *et al*. 2010; Hendry 2016; Snell-Rood *et al*. 2018; Fox *et al*. 2019).

Genetic variation in plasticity should aid population persistence in novel environments when genotypes with plasticity that is at least partially adaptive help to prevent more severe fitness declines (Lande 1988; Lande & Shannon 1996; Bell 2013). These genotypes that are better at maintaining fitness in novel environments can have lower relative fitness in native environments and remain hidden within the native range (Hermisson & Wagner 2004; Angert *et al*. 2008; Brennan *et al*. 2019; Walter *et al*. 2023). Such hidden genetic variation is a critical yet poorly understood source of adaptive potential for populations facing rapid environmental change. By selecting genotypes that differ in their ability to cope with novel environments, and then replicating these genotypes across native and novel environments, it is possible to test how genetic differences in plasticity determine fitness (Chevin *et al*. 2013). Comparing genotypes with higher relative fitness within the native range (HR, ‘Home Range’ genotypes) to genotypes with higher relative fitness in novel environments (AP, ‘Adaptive Potential’ genotypes) provides a powerful framework that harnesses the hidden genetic potential present in populations (**Fig. 1a**; Hermisson & Wagner 2004; Angert *et al*. 2008; Brennan *et al*. 2019; Walter *et al*. 2023).

**Fig. 1.**
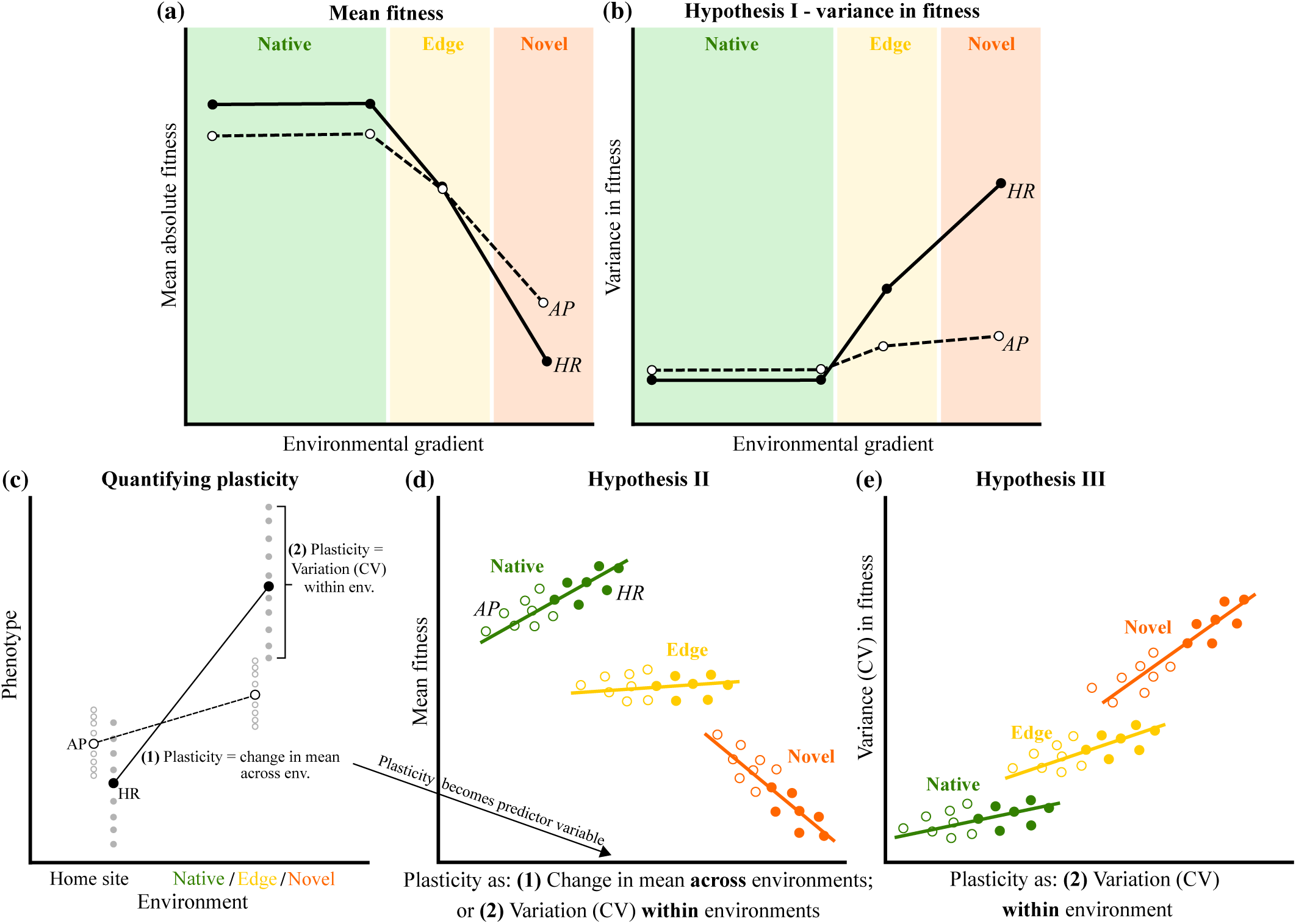
Conceptual framework for testing how genetic variation in plasticity maintains fitness across environmental scales as environments become novel. **(a)** Selection of genotypes that include HR (‘Home Range’; closed circles and solid lines) genotypes that show greater relative fitness in native environments, compared to AP (‘Adaptive Potential’, open circles and dashed lines) genotypes that show greater relative fitness in novel environments. **(b) *Hypothesis I*:** If AP genotypes have partially adaptive plasticity in the novel environment, they would show consistently higher fitness in responses to the novel environment, which would result in lower variance in fitness in the novel environment compared to HR genotypes. **(c)** To connect fitness variation to phenotypic plasticity, we first estimated two plasticity metrics: (1) the change in mean phenotype across environments, and (2) the variation among clones in response to small-scale variation within each environment. One AP genotype (open circles) and one HR genotype (closed circles) is depicted, with small grey circles representing clones within each environment, and large black circles representing the mean of that genotype. Reflecting the original study, we expected AP genotypes to show lower plasticity than HR genotypes. We then tested two hypotheses relating plasticity to fitness: **(d) *Hypothesis II*:** We predicted that the association between plasticity and fitness would change across the elevational gradient. Specifically, if greater phenotypic variation only helps to track changes within native environments, greater plasticity would be favoured within the native range, whereas lower plasticity would be favoured outside the native range. **(e) *Hypothesis III:*** If plasticity in response to fine-scale environmental variation determines fitness, we predicted that in native environments larger plasticity would only be weakly associated with variance in fitness because all genotypes would show similarly high mean fitness. By contrast, at the novel elevation, greater phenotypic changes would incur fitness costs and so greater plasticity would be associated with greater variance in fitness.

The spatial scale at which genetic differences in plasticity determines fitness across ecological gradients is not well understood (De Kort *et al*. 2020; Denney *et al*. 2020). While coarse-scale environmental changes are likely to induce large plastic responses, phenotypic variation in response to fine-scale microenvironmental heterogeneity may generate plasticity of a different magnitude that could be crucial for maintaining fitness across ecological scales (Baythavong 2011; Hamann *et al*. 2016). High replication of HR and AP genotypes across an environmental gradient can then test how genetic differences in fitness arise due to differences in environmental sensitivity both across environments and in response to fine-scale variation within environments. This approach identifies how plasticity varies the phenotype at different ecological scales, and the effect on fitness, to reveal how genetic differences in plasticity could help populations to track environmental variation (e.g., during environmental change or range shifts) or support population persistence as environments become novel (Valladares *et al*. 2007; Valladares *et al*. 2014; Donelson *et al*. 2019; Zettlemoyer 2023; Lewin *et al*. 2024).

Three hypotheses need to be tested to understand how genetic differences in plasticity determines fitness across native and novel environments. ***Hypothesis I:*** Adaptive plasticity reduces variance in fitness because all genotypes make the same adjustments to maintain fitness (Richards *et al*. 2006; Ghalambor *et al*. 2007). Beneficial plasticity in AP genotypes will then mean they consistently adjust their phenotype to better cope with novel environments, and so will show lower variance in fitness at novel environments compared to HR genotypes (**Fig. 1b**). Conversely, HR genotypes should show higher variance in fitness in novel environments if they produce a wider variety of phenotypes that reduce mean fitness. ***Hypotheses II-III*** compare plasticity in response to fine-scale variation within environments, with coarse changes across environments, to test how environmental scale influences plasticity to determine fitness in native and novel environments (**Fig. 1c**). ***Hypothesis II:*** While selection in fine-scale heterogeneous environments favours the evolution of adaptive plasticity (Baythavong & Stanton 2010; Baythavong 2011), we do not know if plasticity with the same magnitude or direction is favoured across native and novel environments, or whether this changes with environmental scale. Higher plasticity should be favoured at fine and coarse scales in native environments because all phenotypic adjustments will maintain high mean fitness. By contrast, novel environments should favour reduced plasticity at both scales because large phenotypic changes become maladaptive (**Fig. 1d**; DeWitt *et al*. 1998; Murren *et al*. 2015; Snell-Rood *et al*. 2018). ***Hypothesis III:*** All plastic responses to fine-scale variation should maintain high fitness in native environments, but if the same plasticity becomes maladaptive in novel environments, then any variation in phenotype will affect fitness and higher plasticity should increase variance in fitness (**Fig. 1e**).

*Senecio* (Asteraceae) wildflower species that inhabit Mt Etna (Sicily) are a powerful system to test how plasticity is linked to adaptive capacity across elevations under semi-natural field conditions (Walter *et al*. 2020). We focus on *S. chrysanthemifolius* that is native to c.400-1500m elevation on Mt Etna. This species is a self-incompatible, short-lived perennial that relies on generalist insect pollinators (e.g., hoverflies). A closely related species, *S. aethnensis,* occurs on old lava flows at high elevations. In previous transplant experiments, the two *Senecio* species showed adaptation to their contrasting habitats associated with differences in plasticity and genetic variance in leaf traits (Walter *et al*. 2022a; Walter *et al*. 2024).

Here we present a large field experiment conducted in 2020, which significantly extends our 2018 field experiment that transplanted cuttings of 314 genotypes of *S. chrysanthemifolius* on Mt Etna (**Box 1**). We showed that greater adaptive potential at a novel 2000m elevation was associated with genetic variance in plasticity (Walter *et al*. 2023). From the 2018 study, we selected 19 genotypes that showed contrasting fitness responses across elevations (**Box 1**). In 2020, we transplanted numerous clones (n=40 per elevation) of each genotype at four elevations and quantified fitness and phenotype to test the three hypotheses outlined above. Transplanting clones at high replication across the entire elevational gradient provided two benefits. First, it allowed us to test whether different forms of plasticity (magnitude and direction) are favoured within the native range compared to novel elevations, which builds on the original study that focused on plasticity at the novel elevation. Second, we could test how plasticity determines fitness at different ecological scales, both in response to large environmental changes across elevations, and fine-scale variation within elevations.

### Box 1

In the 2018 study, we connected genetic variance in plasticity with fitness at a novel 2000m elevation by transplanting cuttings (clones) of 314 genotypes on Mt Etna (Walter *et al*. 2023). Genotypes were generated by mating randomly among 72 individuals that we sampled from five sites located <5km apart at 526-790m elevation on Mt Etna (**Fig. S1**; Walter *et al*. 2023). The 314 genotypes therefore represent genotypes that could be easily generated in the natural population given this species has wind-dispersed seeds and is insect-pollinated. We found no evidence of local adaptation among the sites, and variation in fitness was distributed relatively evenly among parents from different sites (Walter *et al*. 2023). While mean fitness declined in the novel environment, additive genetic variance in fitness increased threefold, reflecting greater adaptive potential at the novel elevation compared to a native elevation. This increased adaptive potential was associated with genetic differences in plasticity. The contrasting fitness responses of these genotypes provide an exceptional opportunity to test how plasticity mediates fitness at different environmental scales along an ecological gradient. From the 2018 experiment, we chose genotypes based on their change in relative fitness (independent of mean absolute fitness) from the home site (500m) to the novel elevation (2000m). Adaptive Potential (AP) genotypes showed greater relative fitness at the novel elevation, whereas Home Range (HR) genotypes showed greater relative fitness at the native elevation (**Fig. 2a**). In the current study, for each of the chosen 2018 genotypes, we transplanted c.40 cuttings at each of four elevations. We recovered the same patterns of mean fitness as the original study: HR genotypes performed better than AP genotypes at native elevations, and AP genotypes performed better at the novel environment (**Fig. 2b**). Our findings were therefore consistent across years, and including an additional 1000m elevation showed that AP genotypes had consistently lower fitness in the native range.

## MATERIAL AND METHODS

From the 2018 experiment, we selected genotypes with the most contrasting fitness responses between 500-2000m: 10 ‘Adaptive Potential’ (AP) genotypes that displayed higher relative fitness in the novel environment, but lower fitness at the native site, and 9 ‘Home Range’ (HR) genotypes that showed higher fitness at their native site, but lower fitness at the novel elevation (**Fig. 2a**). In 2020, we propagated 160 cuttings of each genotype and transplanted them at four elevations representing two sites within their native range (500m and 1000m), their range edge (1500m) and a novel elevation (2000m). We measured leaf traits and fitness on mature plants.

**Fig. 2.**
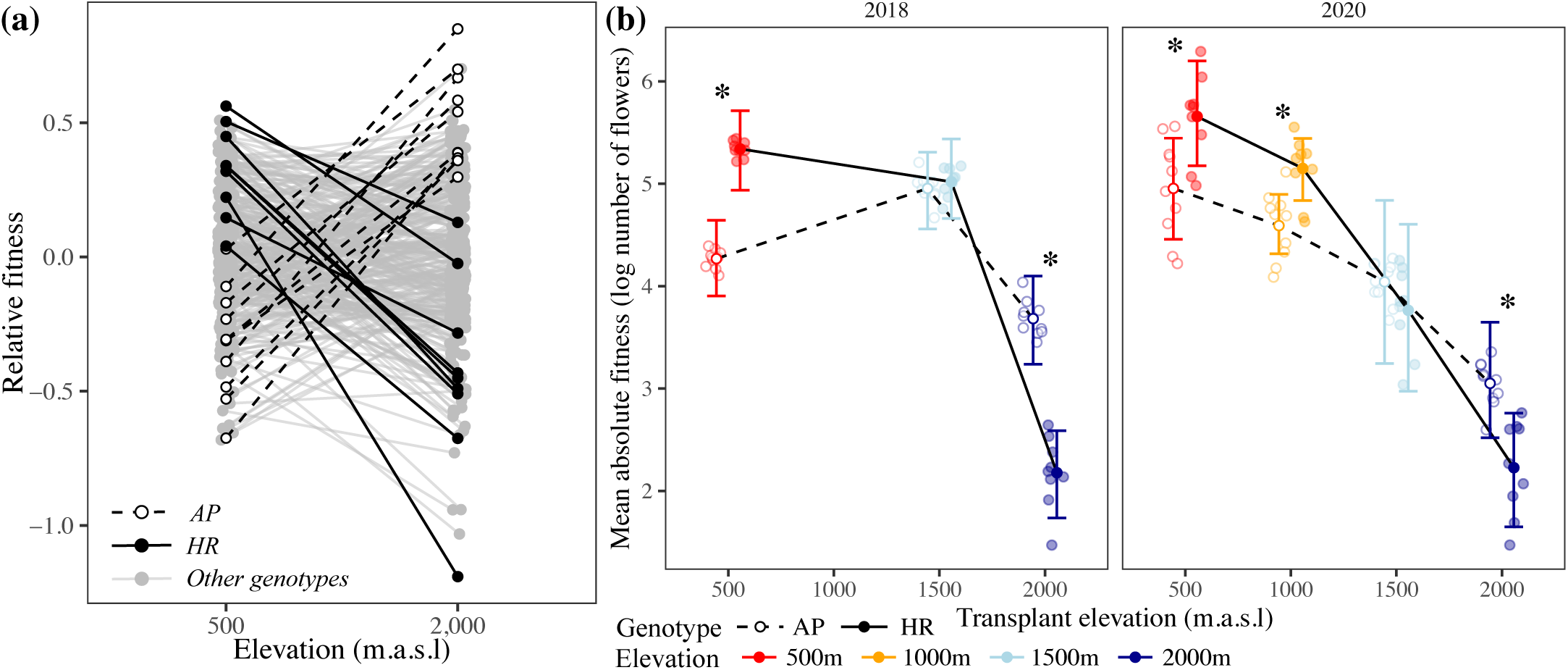
(**a)** Genotypes were chosen from the 2018 experiment based on their change in relative fitness from their home site to the novel 2000m elevation. **(b)** Changes in mean fitness across elevation for the chosen genotypes in the original 2018 study and the current study (2020). Credible intervals represent the 90% Highest Posterior Density interval (HPD) of the mean. Asterisks denote significant differences in mean number of flowers between AP (Adaptive Potential: open circles and dashed lines) and HR (Home Range: closed circles and solid lines) genotypes at each elevation whereby their posterior distributions do not overlap at >90%. Small circles represent the mean for each genotype at each elevation.

### Field transplant

In a greenhouse (Giarre, Italy) in Spring (2020), we propagated 10 clones of each genotype in 14cm diameter pots, randomised their location and let them grow into large plants (c.40cm high). We then removed 6-7 branches from each plant, which we cut into 4cm segments with 2-3 leaf nodes. We dipped the smaller cuttings in rooting hormone (Germon Bew., Der.NAA 0.5%, L.Gobbi, Italy) and placed them in an 84-cell tray containing an equal mix of compressed coconut coir and perlite. We kept trays on benches covered with plastic for three weeks to maintain high humidity and promote root formation.

Rooted cuttings of each genotype were transplanted in early summer (29-30^th^ June) to four elevations 500-2000m on the south-eastern slope of Mt Etna, including a 500m site in a garden among fruit trees, 1000m site in an abandoned vineyard, 1500m site in an abandoned apple orchard, and 2000m site on a lava flow from 1983 with pine trees present (Walter *et al*. 2022a). Higher elevations experience consistently colder temperatures (**Fig. S2**), and soil across the elevation gradient changes from silty sand at elevations 500-1500m to volcanic soil at 2000m (Walter *et al*. 2022a). At each elevation, we randomised the 40 cuttings/genotype into four spatially distinct experimental blocks (*n*=190 plants/block; *n*=760 plants/elevation; N=3040 plants), within an area of c.5000m^2^. At each block, we cleared vegetation and debris, turned the soil 30cm deep, and then planted the cuttings 30cm apart in a grid of 7×29 plants (**Fig. S3**). We irrigated the cuttings daily for three weeks so they could establish, and then reduced irrigation to only the hottest days to prevent high mortality.

### Data collection

We took data for each plant c.4 months after the initial transplant (13-23^rd^ October). As our proxy for fitness, we collected all flowerheads produced by each plant in paper bags, which we counted in the laboratory. This trait is routinely used to estimate fitness in short-lived perennials (e.g., Gross *et al*. 2004; Pujol *et al*. 2014), including in our previous experiment that showed a close association with seed production (Walter *et al*. 2023). To measure five ecologically important leaf traits, we sampled 3-4 fully expanded young leaves from each plant, which we weighed and then scanned to quantify morphology using the program *Lamina* (Bylesjo *et al*. 2008). We used three leaf traits to represent leaf size, shape and investment: leaf area (mm^2^), perimeter (mm) and the number of indentations (count). To calculate the density of leaf indents, we standardized the number of indentations by the perimeter. Using leaf weight, we estimated Specific Leaf Area 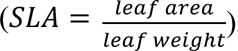, where greater values represent larger leaves per unit mass. With a Dualex instrument (Force-A, France), we measured chlorophyll and flavonol pigment content (light absorbance units) for two leaves per plant. Flavonols are secondary metabolites produced under stressful abiotic (e.g. temperature) and biotic (e.g. herbivore) conditions (Mierziak *et al*. 2014). We chose leaf morphology and investment traits because they show strong plasticity in response to elevation, and are associated with reproductive fitness in *Senecio* (Brennan *et al*. 2009; Walter *et al*. 2023; Walter *et al*. 2024), and other plants (Dudley 1996; Ackerly *et al*. 2000; Van Kleunen & Fischer 2005; Gianoli & Saldaña 2013; Damián *et al*. 2020).

### Hypothesis I: Statistical analyses of fitness

All analyses were conducted in R (v.4.3.2; R Core Team 2024). To quantify mean fitness across elevation, we used *MCMCglmm* (Hadfield 2010) to apply

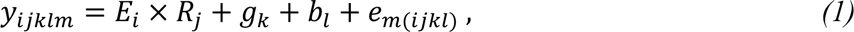

where the interaction between fixed effects of genotype class (𝑅_j_; AP vs HR) and elevation (𝐸_𝑖_) quantifies whether AP and HR genotypes show different fitness responses to elevation. Genotype (𝑔_𝑘_) and experimental block (𝑏_𝑙_) are random effects, and 𝑒_(𝑖j𝑘𝑙)_is the residual. For each random effect, we specified unstructured matrices to estimate variances at each elevation. The number of flowers was the poisson-distributed fitness response variable (𝑦_𝑖j𝑘𝑙𝑚_). Equation 1 yielded the posterior distribution of mean fitness at each elevation. We then applied equation 1 on AP and HR genotypes separately to test whether among-genotype and among-clone (within genotypes, i.e., residual) variance differed between AP and HR genotypes across elevations (*Hypothesis I*).

### Statistical analyses of phenotype and calculation of plasticity

To test whether AP and HR genotypes showed differences in phenotype across elevations, we used *glmmTMB* (Brooks *et al*. 2017) to apply equation 1, but included the five leaf traits as univariate response variables. We used type-III ANOVA (Fox & Weisberg 2019) to test for significant 𝐸_𝑖_ × 𝑅_j_ interactions, which indicate genotypic differences in plasticity to elevation. We then used *emmeans* (Lenth 2019) to obtain marginal means for each genotype and calculate plasticity using

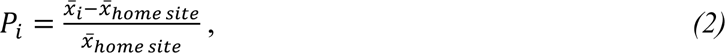

where plasticity (𝑃_𝑖_) for each genotype is the difference in mean between the home site and the *i*th elevation, standardised by the home (500m) site mean (Valladares *et al*. 2006). This captures plasticity as the elevational change in magnitude and direction (negative values reflect a trait decrease) of the phenotype relative to the home site (Anderson *et al*. 2021).

To estimate plasticity within elevations, we calculated the coefficient of variation (CV) for each genotype (and each leaf trait separately) using

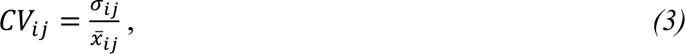

where 𝜎_𝑖j_ and 𝑥̅_𝑖j_ represent the standard deviation and mean, respectively, for the *i*th genotype transplanted at the *j*th elevation. Equation 3 therefore captures plasticity as the among-clone (within-genotype) variance including differences among blocks at each elevation (Hill & Mulder 2010). This is an appropriate use of CV as we are comparing differences between AP and HR genotypes randomised into the same experimental blocks, and because we do not estimate CV across elevations (Pélabon *et al*. 2020). We removed one AP genotype with low replication (<15 clones/elevation) to avoid an imprecise estimate of variance relative to the other genotypes.

### Hypotheses II-III: Connecting plasticity with fitness

In the following analyses, we pooled HR and AP genotypes to use their combined variation in plasticity and phenotype to test how associations with fitness change across elevations. First, to test phenotype-fitness associations, we estimated phenotypic and genotypic selection. We divided each trait by its mean and tested for elevational changes in selection using *glmmTMB* to apply

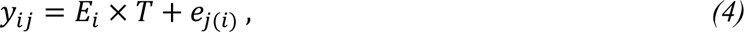

where 𝐸_𝑖_ represents the *i*th elevation and 𝑇 a leaf trait. We included fitness as the response variable (𝑦_𝑖j_) and 𝑒_j(𝑖)_are the residuals. Significant 𝐸_𝑖_ × 𝑇 interactions provide evidence that associations between the trait and fitness changed across elevations. We used the same approach to estimate genotypic selection using genotype means at each elevation (Rausher 1992).

To test whether plasticity changed its association with fitness across elevation, we used equation 4 with plasticity as: (1) the change in trait mean across elevation, and (2) the amount of variation (CV) within elevations. A significant 𝐸_𝑖_ × 𝑇 would provide evidence that the association between plasticity and fitness changed across elevation for that leaf trait. We used the *emtrends* function to test for significant regression slopes.

## RESULTS

### Hypothesis I: Variance in fitness changes across elevations differently for AP and HR genotypes

From low to high elevations, HR genotypes showed a significant 2-3-fold increase in among-genotype variance in fitness. By contrast, AP genotypes showed a significant 3-fold decrease in among-genotype variance in fitness as environments became novel (**Fig. 3a**). In addition, at 1500m and 2000m, HR genotypes showed c.5 times greater among-genotype variation in fitness than AP genotypes (**Fig. 3a**). We found the same patterns for the among-clone (within genotype) variance in fitness: HR genotypes showed an increase and AP genotypes a decrease, in among-clone variance at higher elevations (**Fig. 3b**). Therefore, consistent with *Hypothesis I*, HR genotypes showed (1) greater variance in fitness (within and among genotypes) than AP genotypes at novel elevations, and (2) an increase in genetic variance in fitness at higher elevations. Contrary to predictions, AP genotypes showed a reduction (rather than a slight increase) in variance in fitness at higher elevations.

**Fig. 3.**
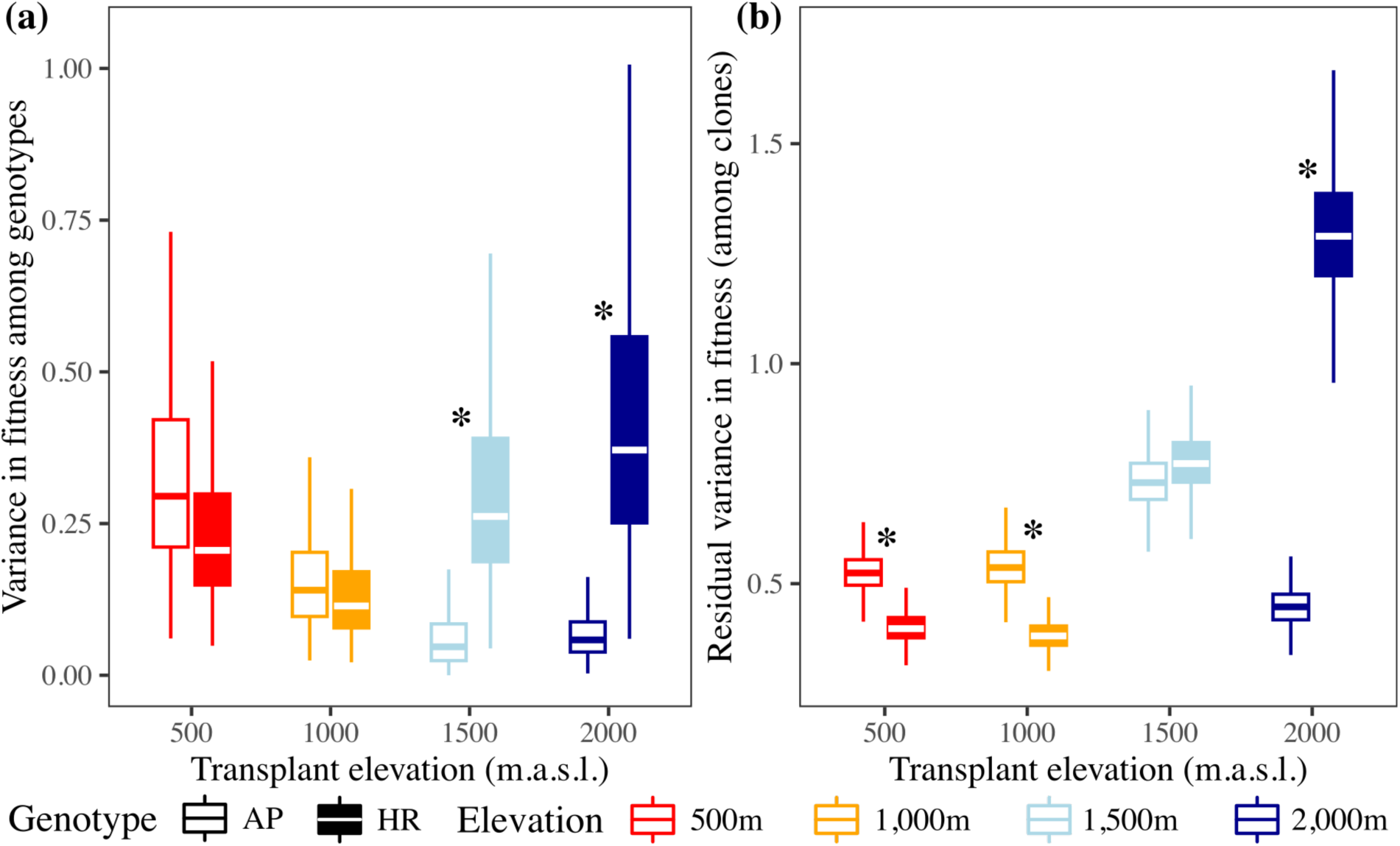
*Hypothesis I:* Fitness variance among genotypes and among clones (within genotypes) changed across elevation differently for AP and HR genotypes, with the greatest difference in variation between the genotypes emerging at the novel elevation. Boxplots represent the posterior distribution of variance in fitness among: **(a)** genotypes, and **(b)** clones within genotype. Unfilled boxplots represent AP genotypes and filled boxplots HR genotypes. Asterisks denote significant differences where the posterior distributions do not overlap at >90%.

### Genotypic differences in plasticity

For both genotypes, all five leaf traits showed reductions in mean values at higher elevations, except for leaf indentation, which increased (**Fig. 4a**). Flavonol content increased at the edge of the range, but then at 2000m returned to similar values as 500m. AP and HR genotypes showed significant differences in plasticity across elevations (i.e. significant genotype class×elevation interactions) for leaf area and chlorophyll content, and differences in mean phenotype across all elevations for specific leaf area and flavonol content (**Fig. 4a; Table S1**).

**Fig. 4.**
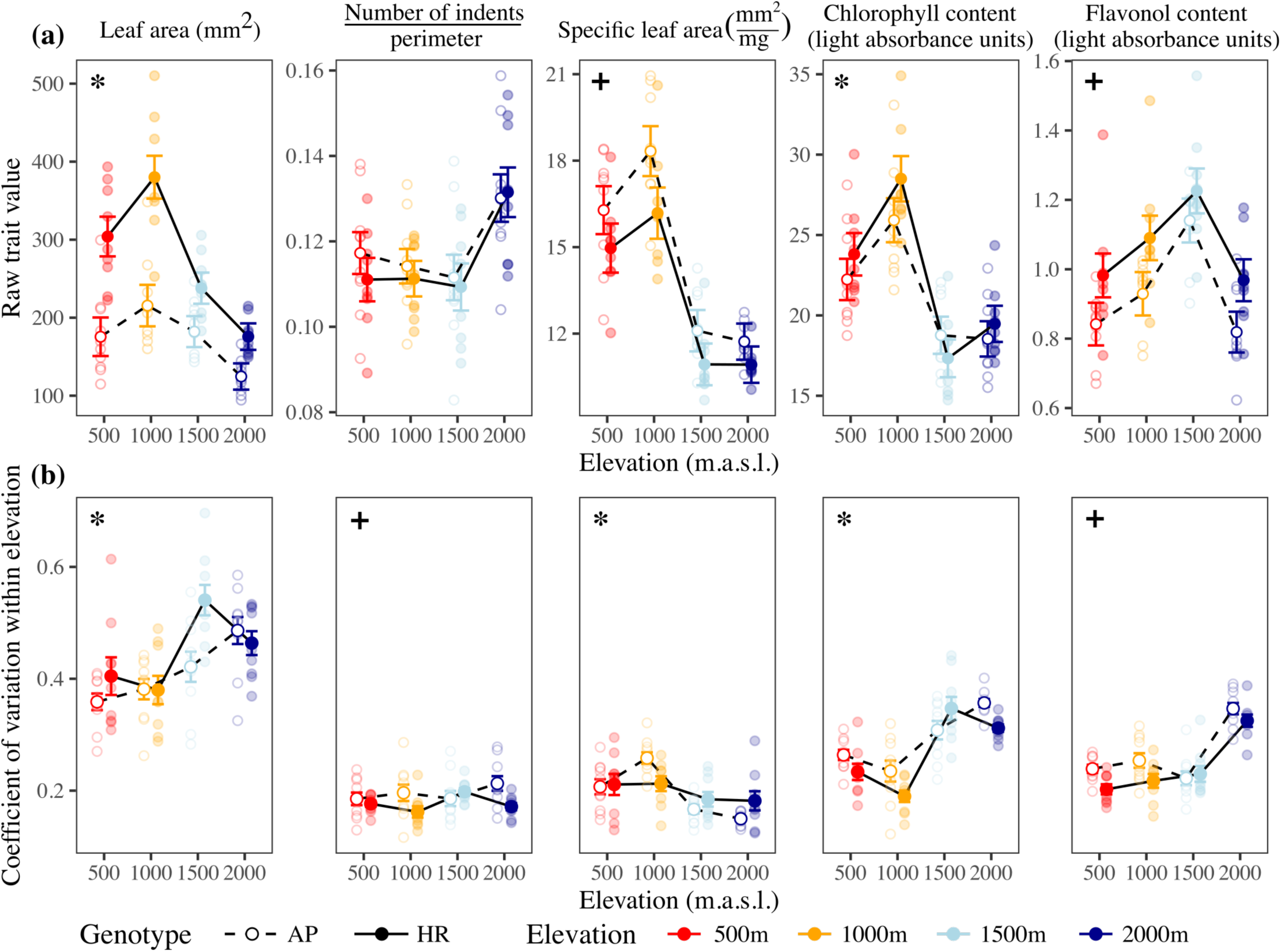
Quantifying plasticity in leaf traits as **(a)** changes in trait means across elevation, and **(b)** and variance (CV) in phenotype within genotypes. AP genotypes are represented by open circles and dashed lines, and HR genotypes by closed circles and solid lines. Larger circles with credible intervals (±1 SE) represent the mean at each elevation, and small circles represent each genotype. Asterisks denote significant elevation×genotype (AP vs. HR) interaction, while plus (+) signs represent no significant interaction but significant differences between AP and HR genotypes. **(a)** For most traits, AP genotypes show smaller changes in phenotype across elevation compared to HR genotypes. **(b)** Most traits show an increase in variation among clones at higher elevations, with AP genotypes often showing greater variance compared to HR genotypes. Summary ANOVA tables are located in **Table S1**.

At each elevation, we found significant differences among the four experimental blocks for most traits and elevations (**Table S2**). However, there was little evidence that AP and HR genotypes responded differently to blocks within elevations (**Table S2**), suggesting that differences between AP and HR were consistent within elevations. Quantifying plasticity as among-clone variance (within genotypes) within elevations, AP and HR genotypes showed differences in plasticity for all traits. For leaf indents and flavonol content, AP genotypes tended to show greater variation among clones (i.e., higher plasticity) than HR genotypes at several elevations (**Fig. 4b**). For leaf area, SLA and chlorophyll content, we found significant differences in the change in CV across elevation (genotype class×elevation interaction; **Fig.4b; Table S1**), suggesting that plasticity as among-clone variation changed across elevation differently for AP and HR genotypes.

### Hypothesis II – Different forms of plasticity were favoured across versus within elevations

All five traits showed significant phenotypic associations between traits and fitness, and, except for the number of leaf indents, higher values of each trait were favoured at all elevations (**Fig. S4; Table S3a**). Genotypic associations between traits and fitness changed significantly across elevation for leaf area, SLA and flavonol content. Higher elevations favoured lower flavonol content, higher SLA, but no association for leaf area. By contrast, lower elevations favoured larger leaves, lower SLA and higher flavonol content (**Fig. 5a; Table S3b**). Genotypes with different trait values were therefore favoured in the native range compared to the novel elevation.

**Fig. 5.**
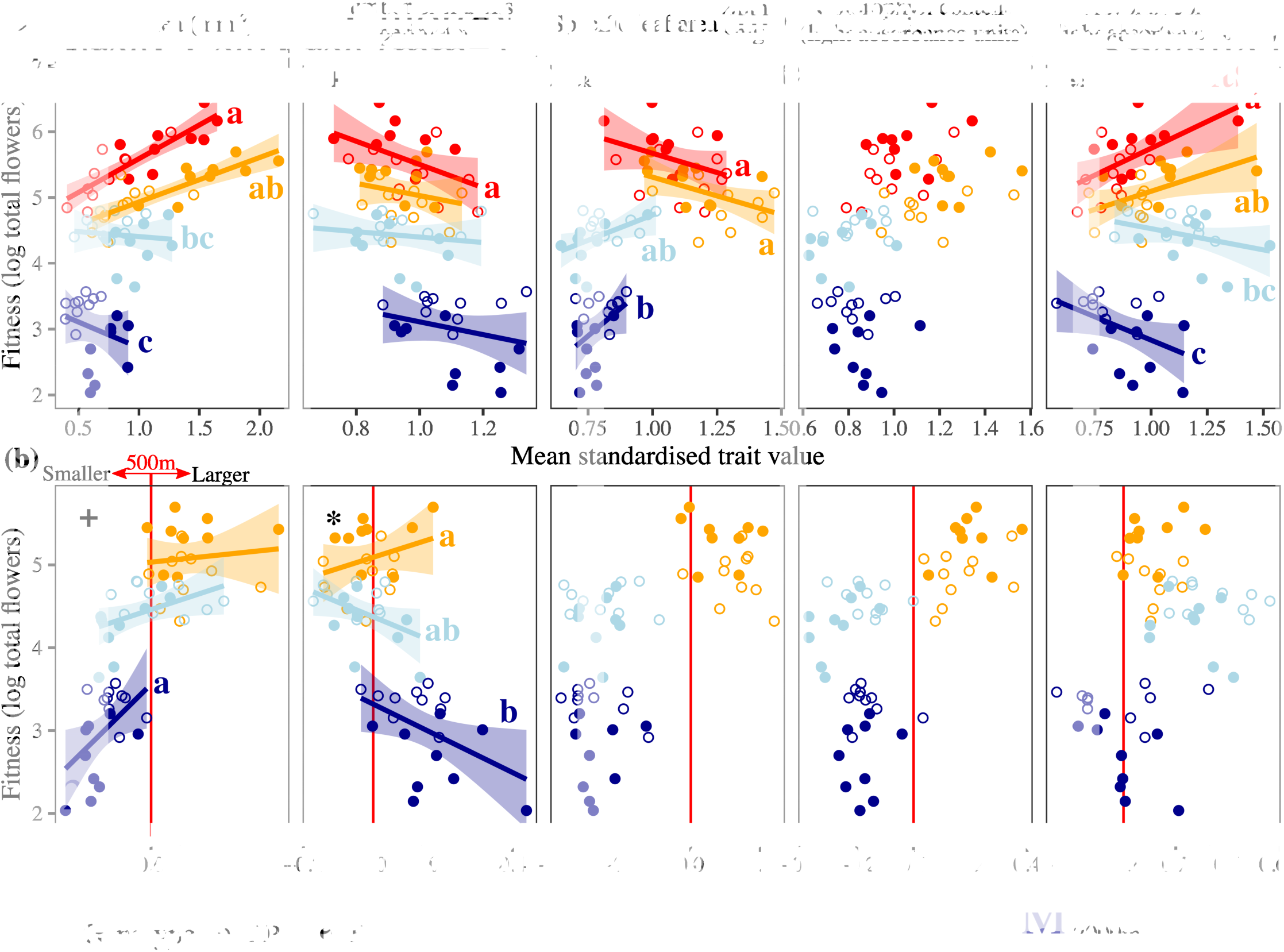
Associations between **(a)** phenotype and **(b)** plasticity with fitness across elevation. AP and HR genotypes are represented by open and closed circles, respectively. Asterisks represent a significant interaction between the predictor variable (trait value or plasticity) and elevation, while addition (+) symbols represent significant regression slopes, but no significant interaction with elevation. Summary ANOVA tables are in **Tables S3-4**. Lines and shaded area (95% confidence intervals) represent the regression, which are omitted for non-significant comparisons. Letters represent significant differences in regression slopes, and panels with a single letter represents a significant slope but no significant differences with other elevations. Regression summary tables are in **Tables S5-6**. **(a)** Genotypic values of each trait on fitnes (phenotypic associations with fitness are located in **Fig. S4**). Three traits (leaf area, specific leaf area and flavonols) showed significant changes in selection on genotypes across elevation. **(b)** *Hypothesis II*: genotypic values of plasticity versus fitness. Plasticity is represented as the change in mean phenotype from the home site (red vertical line) to each other elevation. Positive values represent plasticity as an increase in trait value, and negative values a decrease in trait value, from the home site.

We found support for *Hypothesis II*, that greater plasticity would increase fitness within the native range, but reduce fitness at the novel elevation (**Fig. 1d**). Associations between plasticity and fitness changed significantly across elevation for leaf area and leaf indentation. Only leaf indentation showed a significant elevation×plasticity interaction (F_2,51_=3.6, P=0.035), which meant that larger plastic increases in leaf indentation were favoured within the native range, whereas smaller increases in indentation were favoured at 2000m (**Fig. 5b**). For leaf area, associations between plasticity and fitness were weakly positive at 1000 (slope=0.18±0.4 [1 SE]) and 1500m (slope=0.55±0.3), but strong and significantly positive at 2000m (slope=1.67±0.6, T=2.7, P=0.009; **Fig. 5b**). Because plasticity increased leaf area at low elevations, but decreased leaf area at high elevations, this meant that larger increases (i.e., higher plasticity) in leaf area were associated with slightly higher fitness at 1000-1500m, but smaller decreases (i.e., lower plasticity) in leaf area were strongly favoured at 2000m. Therefore, as predicted, higher fitness at 2000m was associated with lower plasticity in leaf area and indentation.

For each genotype, CV within an elevation represents plasticity as variation in phenotype in response to fine-scale environmental variation. For SLA, higher plasticity within elevations was associated with lower mean fitness, particularly at 1000m and 2000m (slopes: 1000m=-6.1±2.7, T=-2.6, P=0.027; 2000m=-6.1±2.7, T=-2.6, P=0.010; **Fig. 6a**). The association between plasticity (CV) and fitness changed across elevation for leaf indents (plasticity×elevation F_3,64_=3.2, P=0.028) and chlorophyll (plasticity×elevation F_3,64_=7.0, P<0.001). However, our results contradicted *Hypothesis II:* greater variance in phenotype (i.e., greater plasticity) was often associated with lower fitness within the native range, but greater fitness outside the native range (**Fig. 6a**). At the novel elevation, leaf indentation and chlorophyll content showed significant positive associations between plasticity and mean fitness, and only SLA showed the predicted negative trend (**Fig. 6a**).

**Fig. 6.**
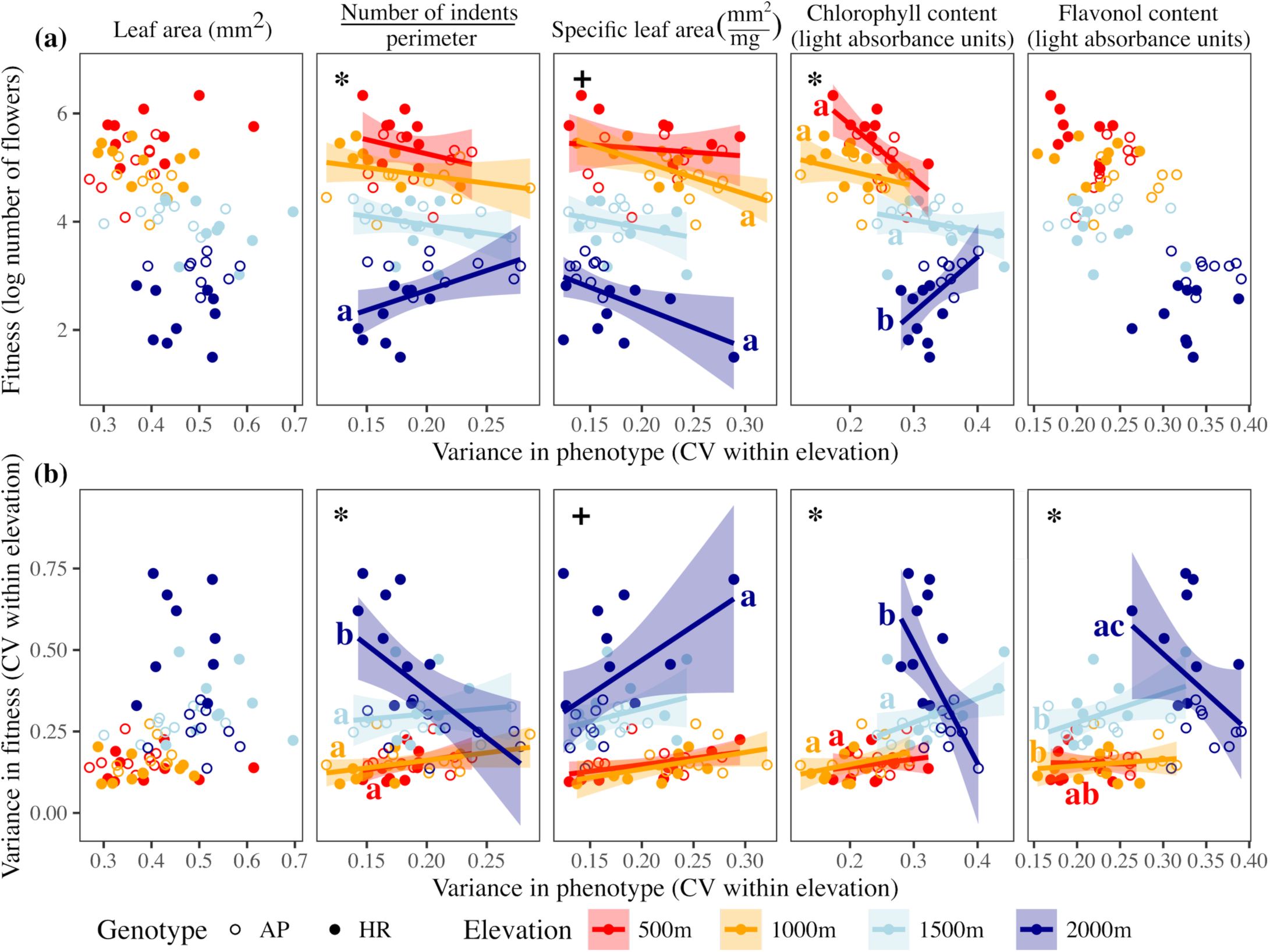
Regression of plasticity as variance (CV) within each genotype at a given elevation against **(a)** mean absolute fitness (*Hypothesis II*), and **(b)** variance (CV) in fitness (*Hypothesis III*). Open circles represent AP genotypes, and closed circles HR genotypes. Asterisks denote significant interaction between plasticity and elevation, and plus (+) signs represent significant slopes but no significant interaction with elevation. Summary ANOVA tables are in **Table S4**. Lines and shaded area (95% confidence intervals) represent the regression, which are omitted for non-significant comparisons. Letters represent significant differences in regression slopes, and panels with a single letter represents a significant slope but no significant differences with other elevations. Regression summary tables are in **Tables S5-6**.

### Hypothesis III – Greater plasticity was associated with lower variance in fitness at the novel elevation

We predicted that higher among-clone variation in phenotype (i.e., greater plasticity) within a genotype would be associated with low variation in fitness within the range, but high variation in fitness at the novel environment (**Fig. 1f**). Supporting *Hypothesis III*, four (of five) traits showed significant associations between CV in phenotype and variance in fitness (**Fig. 6b**). As predicted, we found a weak positive association between CV in phenotype and variance in fitness at elevations within the native range, suggesting that plasticity maintained similarly high fitness for all clones and genotypes. Also as predicted, three traits showed significant changes in the association between CV in phenotype and variance in fitness across elevations (plasticity×elevation for leaf indentation F_3,64_=7.6, P<0.001; chlorophyll F_3,64_=10.8, P<0.001; and flavonols F_3,64_=4.0, P=0.011; **Fig. 6b**). However, contrary to our predictions, these traits showed a strong negative association with variance in fitness at the novel elevation, suggesting that greater plasticity reduced variance in fitness. Only SLA showed the predicted significant positive association at the novel elevation (slope=2.1±0.6, T=3.5, P=0.001; **Fig. 6b**).

## DISCUSSION

To test how genetic differences in plasticity influence fitness across an ecological gradient, we transplanted large numbers of clones of 19 genotypes at four elevations on Mt Etna, Sicily. Genotypes were selected from a previous experiment based on their contrasting fitness response to elevation (Walter *et al*. 2023): Home Range (HR) genotypes showed higher relative fitness at native elevations, but lower relative fitness than Adaptive Potential (AP) genotypes at the novel 2000m elevation (**Box 1**). Supporting *Hypothesis I*, HR genotypes showed increased variation in fitness at higher elevations, whereas AP genotypes showed a decrease (**Fig. 3**). At the novel environment, AP genotypes therefore coped better and showed more consistent responses, whereas HR genotypes produced a variety of responses that resulted in lower mean fitness.

Supporting *Hypothesis II*, genotypic differences in plasticity were associated with fitness. For plasticity across elevations, slightly higher plasticity in leaf area and indentation was favoured within the native range, whereas lower plasticity was favoured at the novel elevation (**Fig. 5b**). However, plasticity as variation within elevations showed a contrasting result: lower plasticity was often favoured within native elevations, whereas higher plasticity was favoured within the novel elevation (**Fig. 6a**). Consistent with *Hypothesis III*, plasticity within native elevations was weakly associated with variance in fitness, suggesting that phenotypic changes in native environments maintained high mean fitness. However, contrary to *Hypothesis III*, higher plasticity was often associated with lower variance in fitness at the novel elevation (**Fig. 6b**).

Our results provide two important insights for understanding how genetic variation in plasticity determines fitness across environments. First, we provide some of the first evidence that genotypes with different forms (magnitude and direction) of plasticity are favoured within their native range compared to novel environments. Second, the association between plasticity and fitness changed in response to fine- vs coarse- scale environmental variation, and was trait-dependent. Higher fitness at novel elevations was generally associated with smaller phenotypic adjustments across elevations, but larger adjustments within elevations.

### Implications for predicting population persistence under global change

We provide some of the strongest evidence that where populations face novel environments, adaptive plasticity fails to maintain high mean fitness, and genetic variation in plasticity will be important for population persistence. By focussing on genotypes with contrasting fitness responses to elevation, we show that selection on plasticity changes across elevation, and that genotypes that differ in plasticity track native and novel environments differently, which also depends on the spatial scale of environmental variation. Moving beyond demonstrating genetic variance in plasticity, we reveal how hidden genetic variation in plasticity could allow new forms of adaptive plasticity to evolve (Lande 2009; Usui *et al*. 2023), help populations to maintain fitness and persist as environments change (Chevin & Hoffmann 2017), or help them shift their geographical range in response to global change (Valladares *et al*. 2007; Valladares *et al*. 2014; Donelson *et al*. 2019; Zettlemoyer 2023; Lewin *et al*. 2024). The frequency and genetic basis of such genotypes, as well as their absolute fitness in novel environments, will determine their potential to aid population persistence under environmental change.

Predicting the resilience of ecological communities requires expanding our framework to multiple species and assaying genotypes from across a metapopulation in conditions predicted under climate change. This would test whether AP genotypes are broadly common across populations and species, or whether they are only present in species with large populations that experience greater environmental heterogeneity. Encouragingly, a field experiment with both Etnean *Senecio* species showed increased genetic variance in seedling survival at both novel (high and low) elevations (Walter *et al*. 2022b), suggesting that AP genotypes are present at early life history stages, and in species from contrasting environments that experienced different novel environments. Testing for AP genotypes in species showing range stasis versus expansion or shifts (or species with different sized distributions) would confirm whether AP genotypes emerge as a broader phenomenon across different ecological contexts for coping with environmental change. Identifying AP genotypes is challenging but would benefit conservation efforts to increase resilience in species from vulnerable ecosystems.

### Genetic variation for population persistence under environmental change

AP genotypes could be maintained in populations through interannual or seasonal variation. Environmental variation that creates fluctuating selection would then maintain genetic variation in plasticity that reduces vulnerability to environmental change (Gillespie & Turelli 1989; Svardal *et al*. 2011; Wittmann *et al*. 2017). AP genotypes could then represent ‘generalist’ genotypes that have lower arithmetic fitness within their native range, but high geometric fitness across a broader range of spatial or temporal environments (Gillespie 1974; Childs *et al*. 2010; Svardal *et al*. 2011; Tufto 2015; Draghi 2023). This would suggest that AP genotypes are ‘bet-hedging’ genotypes that help to buffer large environmental variation by reducing the fitness costs to the population as environments change rapidly and/or unpredictably (Gillespie 1974; Childs *et al*. 2010; Simons 2011; Bond *et al*. 2021; Draghi 2023). Alternatively, differences between AP and HR genotypes in their plasticity and fitness across elevations could be created by different alleles associated with local adaptation to high and low elevations within their native range. Local adaptation could then create and maintain AP genotypes (Lind & Johansson 2007). However, we previously found little evidence of local adaptation among sampling sites, and so further work is needed to understand whether local adaptation generates AP genotypes. Testing whether AP genotypes are broadly beneficial across many novel environments, including warmer conditions at lower elevations, or whether their fitness benefits are specific to high elevations would directly test whether local adaptation generates AP genotypes.

### Contrasting patterns of plasticity in response to coarse versus fine-scale environmental variation

Our results suggest plasticity of different magnitudes was favoured at different ecological scales. Across elevations, higher plasticity tended to be favoured within the native range, whereas lower plasticity was favoured at the novel elevation. This suggests that phenotypic robustness maintains higher fitness in novel environments by minimising large irreversible and costly phenotypic changes (Velotta & Cheviron 2018; Hoffmann & Bridle 2022). By contrast, for plasticity within elevations, lower plasticity was generally favoured within the native range, and higher plasticity favoured at the novel elevation. This supports the idea that phenotypic responses to fine-scale environmental variation are under genetic control and contribute to the genetic basis of adaptive plasticity (Baythavong 2011; Prentice *et al*. 2020). However, the association between plasticity and fitness across ecological scales changed depending on the trait. While leaf area and indentation showed the strongest association between plasticity across elevations and fitness, plasticity in chlorophyll content within elevations showed the strongest association with fitness. Plasticity in different traits could therefore mediate fitness at different ecological scales as environments become novel (Chevin *et al*. 2010).

Identifying the molecular mechanisms that determine how AP genotypes track novel variation at fine and coarse scales will help to predict population persistence under global change. Previously, AP genotypes showed larger changes in gene expression across elevations than HR genotypes (Walter *et al*. 2023), suggesting that AP genotypes maintain higher fitness at the novel elevation through larger changes in gene expression across elevations and more fine-scale phenotypic adjustments within elevation. Fine-scale adjustments could be less important within the native range where large phenotypic responses to reliable environmental cues maintain fitness, whereas the same plasticity becomes costly in response to unreliable cues in novel environments that favour phenotypic stability and a fine-tuning of phenotype. Such contrasting patterns of plasticity could be created by differences in the response to large climatic changes across elevation compared to variation in biotic (e.g., soil biota) conditions within elevations (Paquette & Hargreaves 2021).

### Idiosyncrasies that could determine the generality of AP genotypes

In contrast with our result (plasticity of different magnitudes favoured at coarse vs fine scales), plasticity in tree growth was positively correlated across micro- and macro-environmental variation (de la Mata & Zas 2023). We used flower output as our fitness metric, which may not reflect variation in growth or survival, and so future experiments should test whether AP genotypes emerge using other performance measures, such as biomass, which could be valuable to conservation efforts seeking to increase resilience in long-lived species. Furthermore, the autocorrelation of changes in edaphic (and other environmental variables) along with temperature across elevation meant we could not isolate thermal plasticity relevant to predicting responses to global change. Future work should therefore separate temperature from other environmental variables to identify whether, and at what scales, AP genotypes could aid population persistence to global change.

### Conclusions

We show how hidden genetic variation in plasticity could aid population persistence in novel environments by better tracking coarse and fine variation in the environment that reduces the severity of fitness declines. We also demonstrate that different forms of plasticity are favoured across a natural ecological gradient from native to novel environments. To predict the potential for AP genotypes to aid population persistence under global change, future experiments should focus on their frequency within and across species, and the generality of their fitness benefits across novel environments.

## Acknowledgements

We thank Matt Hall for providing comments on an earlier version, and we thank two anonymous reviewers whose insightful comments helped improve this manuscript. We are very grateful to Piante Faro (Giarre, Italy) for providing us with glasshouse facilities that made this study possible, and we thank Giuseppe Riggio for generously providing us access to the 1000m field site. This work was supported by joint NERC grants NE/P001793/1 and NE/P002145/1 awarded to JB and SH. GW was supported by Australian Research Council fellowships DE200101019 and FT240100466. DT was funded by PON “Ricerca e Innovazione” 2014-2020 - Azione IV.5 “Dottorati su tematiche Green”.

