## Supplementary material for "Genetic differences in plasticity across environmental scales determine fitness along an ecological gradient"

**Contents**

**Fig. S1** Locations of parental genotypes from the natural population

**Fig. S2** Climate comparison of 2018 and 2020

**Fig. S3** Photos of experimental plants

**Fig. S4** Phenotypic selection

**Table S1** ANOVA summary tables for plastic changes in leaf phenotypes across and within elevations

**Table S2** Tests of variation among blocks at each elevation for AP and HR genotypes

**Table S3** ANOVA summary tables for selection on leaf traits

**Table S4** ANOVA summary tables for hypotheses II & III – associations between plasticity and fitness

**Table S5** Slope estimates for regressions in **Figs 5-6**.

**Table S6** Pairwise tests of slope differences between elevations in **Figs 5-6**.

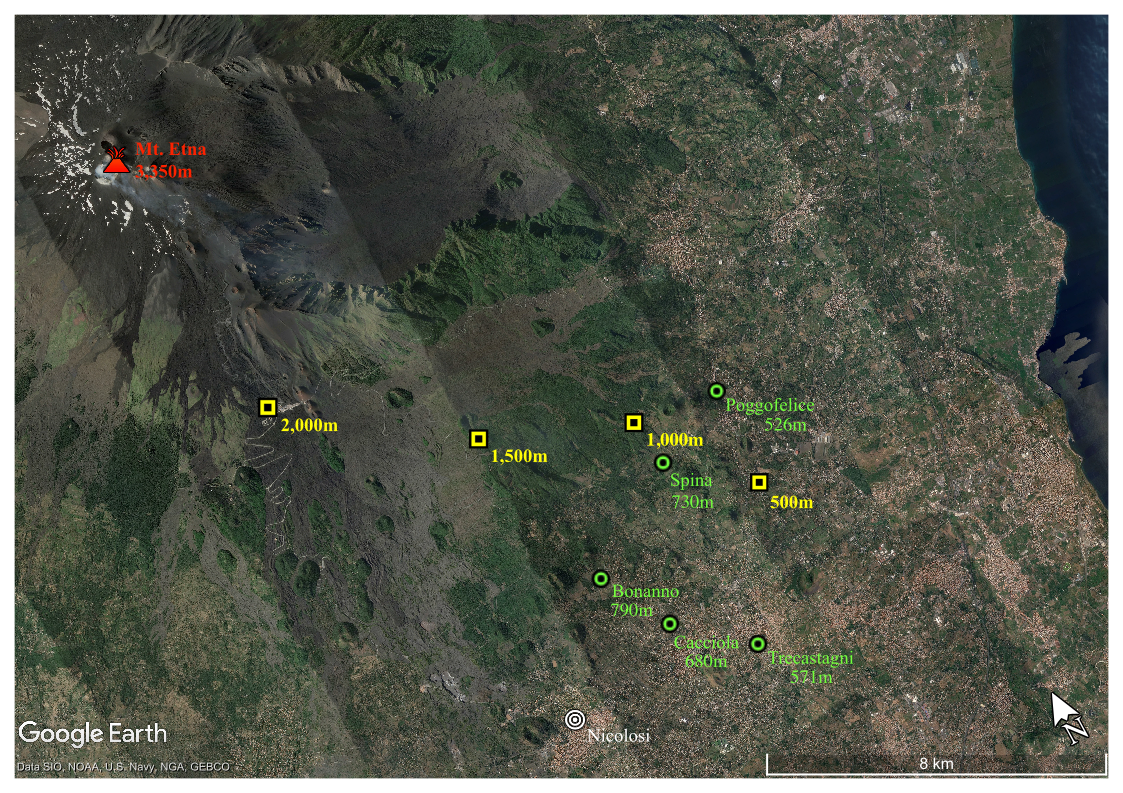

**Fig. S1** Map of transplant sites (yellow and black squares) on Mt Etna, and locations of samples from where parental genotypes were taken in the natural population (green and black circles). Numbers represent elevation as metres above sea level.

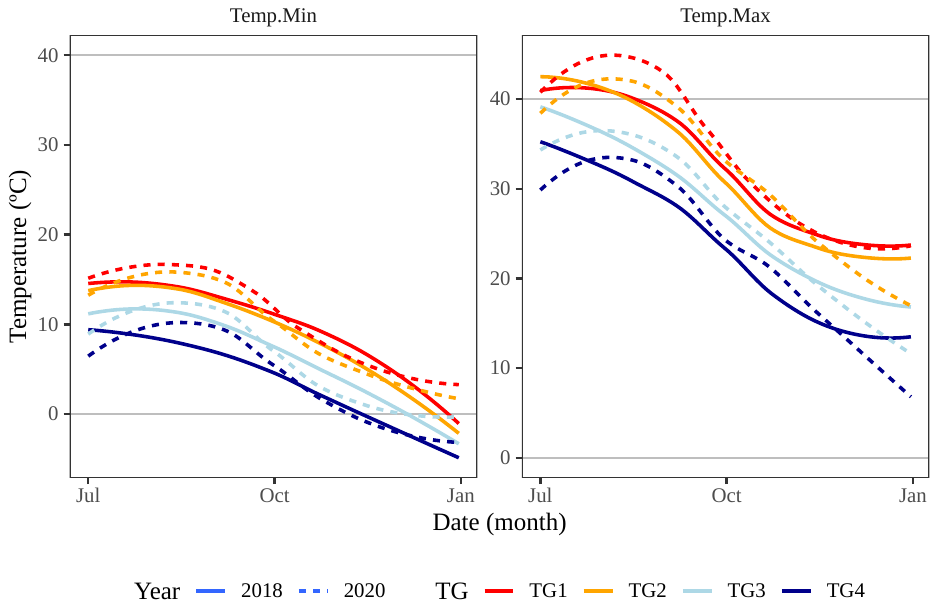

**Fig. S2** Minimum and maximum daily temperatures recorded during both experiments (2018 in solid lines and 2020 in broken lines).

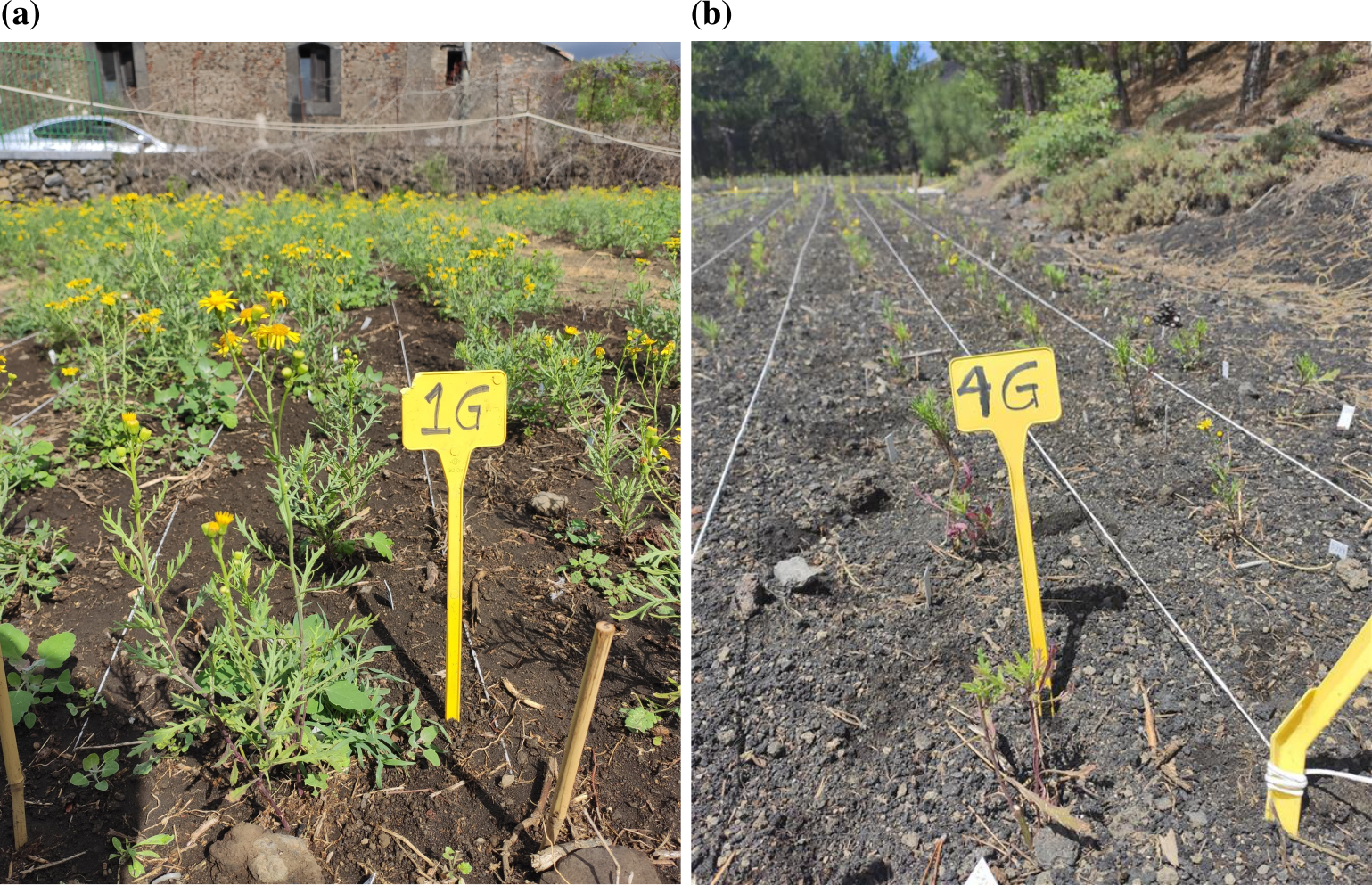

**Fig. S3** Images of experimental blocks in the field at **(a)** 500m and **(b)** 2000m 1 month after the initial transplant.

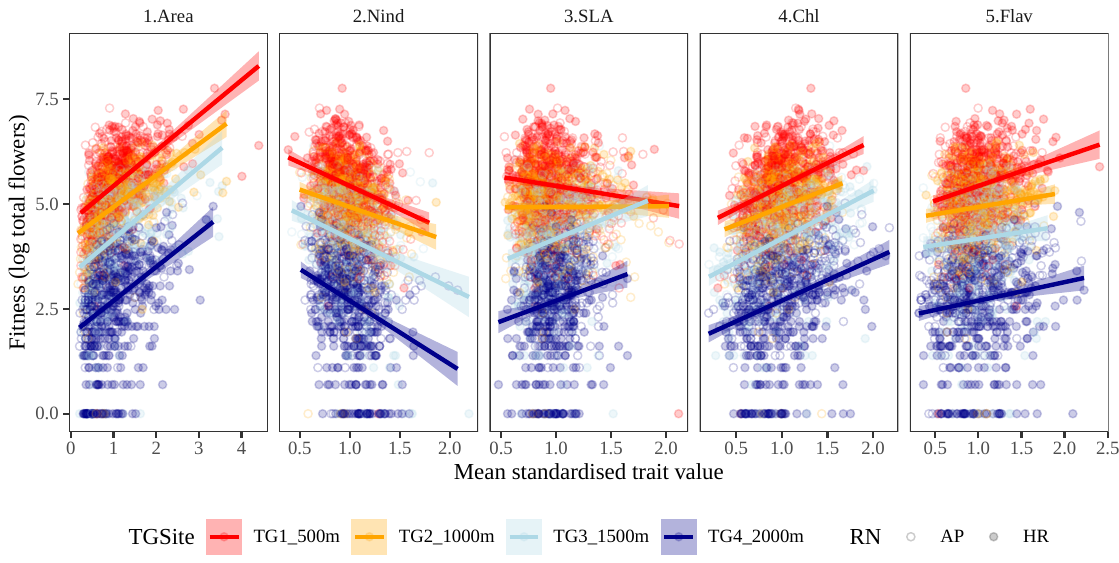

**Fig. S4** Phenotypic selection for all five traits at the four transplant elevations. Circles represent individual plants.

| **Fig. 4a** - Genotypic differences in plasticity across elevation | | | | |  | **Fig. 4b** - Genotypic differences in plasticity within elevation (CV) | | | | | |
| --- | --- | --- | --- | --- | --- | --- | --- | --- | --- | --- | --- |
| **Trait** | **Parameter** | **Chisq** | **Df** | **P-value** |  | **Trait** | **Parameter** | **SS** | **Df** | **F-value** | **P-value** |
| Area | Intercept | 336.43 | 1 | <0.001 |  | Area | Intercept | 13.66 | 1 | 2774.42 | <0.001 |
|  | Elevation | 39.64 | 3 | <0.001 |  |  | Elevation | 0.19 | 3 | 12.62 | <0.001 |
|  | Class | 24.66 | 1 | <0.001 |  |  | Class | 0.01 | 1 | 2.01 | 0.161 |
|  | Elevation × Class | 19.63 | 3 | **<0.001** |  |  | Elevation × Class | 0.06 | 3 | 3.8 | **0.014** |
| Nind | Intercept | 1337.7 | 1 | <0.001 |  |  | Residuals | 0.32 | 64 |  |  |
|  | Elevation | 44.15 | 3 | <0.001 |  | Nind | Intercept | 2.52 | 1 | 2165.39 | <0.001 |
|  | Class | 0.17 | 1 | 0.683 |  |  | Elevation | 0 | 3 | 0.69 | 0.562 |
|  | Elevation × Class | 2.11 | 3 | 0.549 |  |  | Class | 0.01 | 1 | 6.45 | **0.014** |
| SLA | Intercept | 1066.6 | 1 | <0.001 |  |  | Elevation × Class | 0.01 | 3 | 2.05 | 0.116 |
|  | Elevation | 65.83 | 3 | <0.001 |  |  | Residuals | 0.07 | 64 |  |  |
|  | Class | 4.56 | 1 | **0.033** |  | SLA | Intercept | 2.78 | 1 | 1736.87 | <0.001 |
|  | Elevation × Class | 4.36 | 3 | 0.225 |  |  | Elevation | 0.05 | 3 | 11.03 | <0.001 |
| Chl | Intercept | 855.79 | 1 | <0.001 |  |  | Class | 0 | 1 | 0.05 | 0.825 |
|  | Elevation | 72.07 | 3 | <0.001 |  |  | Elevation × Class | 0.01 | 3 | 2.98 | **0.038** |
|  | Class | 0.5 | 1 | 0.480 |  |  | Residuals | 0.1 | 64 |  |  |
|  | Elevation × Class | 14.05 | 3 | **0.003** |  | Chl | Intercept | 5.65 | 1 | 3436.64 | <0.001 |
| Flav | Intercept | 765.39 | 1 | <0.001 |  |  | Elevation | 0.22 | 3 | 44.07 | <0.001 |
|  | Elevation | 28.63 | 3 | <0.001 |  |  | Class | 0.01 | 1 | 4.05 | **0.049** |
|  | Class | 4.92 | 1 | **0.027** |  |  | Elevation × Class | 0.01 | 3 | 3.02 | **0.036** |
|  | Elevation × Class | 3.33 | 3 | 0.344 |  |  | Residuals | 0.11 | 64 |  |  |
|  |  |  |  |  |  | Flav | Intercept | 4.72 | 1 | 3879.08 | <0.001 |
|  |  |  |  |  |  |  | Elevation | 0.17 | 3 | 45.66 | <0.001 |
|  |  |  |  |  |  |  | Class | 0.01 | 1 | 9.18 | **0.004** |
|  |  |  |  |  |  |  | Elevation × Class | 0.01 | 3 | 1.59 | 0.200 |
|  |  |  |  |  |  |  | Residuals | 0.08 | 64 |  |  |

**Table S1** ANOVA summary tables for plasticity in leaf traits for **(a)** plasticity across elevations, and **(b)** plasticity as phenotypic variation within elevation (CV), and how CV changes across elevations for the two genotype classes. Parameters in bold are significant at P<0.05 to aid interpretation, focusing on the genotypes class (AP vs HR) and the interaction with elevation.

**Table S2** ANOVA summary tables for testing differences among genotype classes and experimental blocks at each elevation. Parameters in bold are significant at P<0.05. Class represents the genotype classes (AP vs HR). Only leaf area at 500m, indents at 500m, SLA at 1000m and 2000m, and flavonols at 1000m showed significant Class×Block interactions. Therefore, for most traits and elevations, the two genotype classes did not differ in their response to variation within elevations. This suggests that plasticity captured by within-elevation variation is not biased by responses to block-specific effects.

|  |  |  | **Leaf area** | | **Indents** | | **SLA** | | **Chlorophyll** | | **Flavonols** | | **Fitness (flowers)** | |
| --- | --- | --- | --- | --- | --- | --- | --- | --- | --- | --- | --- | --- | --- | --- |
| **Elevation** | **Parameter** | **df** | $\boldsymbol{\chi}^{\boldsymbol{2}}$ | **P** | $\boldsymbol{\chi}^{\boldsymbol{2}}$ | **P** | $\boldsymbol{\chi}^{\boldsymbol{2}}$ | **P** | $\boldsymbol{\chi}^{\boldsymbol{2}}$ | **P** | $\boldsymbol{\chi}^{\boldsymbol{2}}$ | **P** | $\boldsymbol{\chi}^{\boldsymbol{2}}$ | **P** |
| 500m | Class | 2 | 317.55 | **<0.001** | 1411.06 | **<0.001** | 1424.3 | **<0.001** | 1036.52 | **<0.001** | 822.67 | **<0.001** | 2765.92 | **<0.001** |
|  | Block | 3 | 14.23 | **0.003** | 24.2 | **<0.001** | 65.35 | **<0.001** | 33.54 | **<0.001** | 24.20 | **<0.001** | 205.68 | **<0.001** |
|  | Class×Block | 3 | 2.07 | 0.559 | 9.87 | **0.02** | 4.07 | 0.254 | 1.79 | 0.617 | 2.53 | 0.469 | 2.27 | 0.519 |
| 1000m | Class | 2 | 389.82 | **<0.001** | 2228.8 | **<0.001** | 1499.81 | **<0.001** | 1235.04 | **<0.001** | 966.19 | **<0.001** | 4759.14 | **<0.001** |
|  | Block | 3 | 3.65 | 0.302 | 2.38 | 0.498 | 88.71 | **<0.001** | 78.51 | **<0.001** | 10.59 | **0.014** | 28.67 | **<0.001** |
|  | Class×Block | 3 | 0.48 | 0.923 | 4.72 | 0.194 | 8.63 | **0.035** | 7.19 | 0.066 | 12.17 | **0.007** | 0.58 | 0.902 |
| 1500m | Class | 2 | 505.58 | **<0.001** | 1066.81 | **<0.001** | 1531.1 | **<0.001** | 981.33 | **<0.001** | 1215.55 | **<0.001** | 3712.13 | **<0.001** |
|  | Block | 3 | 61.52 | **<0.001** | 6.03 | 0.11 | 17.6 | **<0.001** | 2.72 | 0.436 | 39.35 | **<0.001** | 452.91 | **<0.001** |
|  | Class×Block | 3 | 8.62 | **0.035** | 5.23 | 0.156 | 7.21 | 0.066 | 1.47 | 0.690 | 1.70 | 0.638 | 4.17 | 0.244 |
| 2000m | Class | 2 | 615.8 | **<0.001** | 1367.32 | **<0.001** | 4282.32 | **<0.001** | 1285.31 | **<0.001** | 923.91 | **<0.001** | 1514.52 | **<0.001** |
|  | Block | 3 | 95.68 | **<0.001** | 28.39 | **<0.001** | 77.39 | **<0.001** | 26.96 | **<0.001** | 95.26 | **<0.001** | 82.34 | **<0.001** |
|  | Class×Block | 3 | 7.74 | 0.052 | 2.97 | 0.396 | 10.08 | **0.018** | 3.50 | 0.320 | 1.05 | 0.788 | 2.75 | 0.432 |

**Table S3** ANOVA summary tables for selection on **(a)** phenotypic selection, and **(b)** genotype selection (**Fig. 5a**) on leaf traits across elevation. P-values in bold are <0.05 to enhance visualisation of significant main effect of the trait and the interaction with elevation.

| **(a) Phenotypic selection** | | | | |  | **(b) Genotypic selection** | | | | | |
| --- | --- | --- | --- | --- | --- | --- | --- | --- | --- | --- | --- |
| **Trait** | **Parameter** | **Chisq** | **Df** | **P** |  | **Trait** | **Parameter** | **SS** | **Df** | **F** | **P** |
| Area | Intercept | 20330 | 1 | <0.001 |  | Area | Intercept | 104.33 | 1 | 949.64 | <0.001 |
|  | **Area** | 536.61 | 1 | **<0.001** |  |  | Area | 0.15 | 1 | 1.33 | 0.252 |
|  | Elevation | 547.61 | 3 | <0.001 |  |  | Elevation | 0.94 | 3 | 2.87 | 0.043 |
|  | Area × Elevation | 6.02 | 3 | 0.111 |  |  | **Area × Elevation** | 1.78 | 3 | 5.41 | **0.002** |
| Nind | Intercept | 6321.6 | 1 | <0.001 |  |  | Residuals | 7.47 | 68 |  |  |
|  | **Indentation** | 128.12 | 1 | **<0.001** |  | Nind | Intercept | 29.87 | 1 | 188.23 | <0.001 |
|  | Elevation | 96.16 | 3 | <0.001 |  |  | **Indentation** | 0.98 | 1 | 6.17 | **0.015** |
|  | Indentation × Elevation | 3.51 | 3 | 0.319 |  |  | Elevation | 1.59 | 3 | 3.33 | 0.025 |
| SLA | Intercept | 3983.1 | 1 | <0.001 |  |  | Indentation × Elevation | 0.26 | 3 | 0.54 | 0.658 |
|  | SLA | 10.86 | 1 | <0.001 |  |  | Residuals | 10.79 | 68 |  |  |
|  | Elevation | 214.96 | 3 | <0.001 |  | SLA | Intercept | 14.61 | 1 | 100.87 | <0.001 |
|  | **SLA × Elevation** | 42.24 | 3 | **<0.001** |  |  | SLA | 0.26 | 1 | 1.78 | 0.187 |
| Chl | Intercept | 6385.7 | 1 | <0.001 |  |  | Elevation | 4.83 | 3 | 11.11 | <0.001 |
|  | **Chlorophyll** | 211.34 | 1 | **<0.001** |  |  | **SLA** × **Elevation** | 2.08 | 3 | 4.79 | **0.004** |
|  | Elevation | 141.69 | 3 | <0.001 |  |  | Residuals | 9.85 | 68 |  |  |
|  | Chlorophyll × Elevation | 6.61 | 3 | 0.085 |  | Chl | Intercept | 20.29 | 1 | 132.41 | <0.001 |
| Flav | Intercept | 6924.3 | 1 | <0.001 |  |  | Chlorophyll | 0.53 | 1 | 3.43 | 0.068 |
|  | Flavonol | 39.87 | 1 | <0.001 |  |  | Elevation | 0.03 | 3 | 0.06 | 0.979 |
|  | Elevation | 123.78 | 3 | <0.001 |  |  | Chlorophyll × Elevation | 0.71 | 3 | 1.54 | 0.212 |
|  | **Flavonol × Elevation** | 14.09 | 3 | **0.003** |  |  | Residuals | 10.42 | 68 |  |  |
|  |  |  |  |  |  | Flav | Intercept | 34.88 | 1 | 254.66 | <0.001 |
|  |  |  |  |  |  |  | Flavonol | 0.05 | 1 | 0.37 | 0.546 |
|  |  |  |  |  |  |  | Elevation | 0.32 | 3 | 0.78 | 0.511 |
|  |  |  |  |  |  |  | **Flavonol × Elevation** | 2.69 | 3 | 6.54 | **0.001** |
|  |  |  |  |  |  |  | Residuals | 9.31 | 68 |  |  |

**Table S4** ANOVA summary tables for associations between fitness and plasticity. **(a)** *Hypothesis II -* Associations between mean fitness and plasticity across elevations in **Fig. 5b**, **(b)** *Hypothesis II -* Associations between mean fitness and plasticity as variance in phenotype within elevations (CV) in **Fig. 6a**, and **(c)** *Hypothesis III -* Associations between variance in fitness and plasticity as CV within elevations in **Fig. 6b**, and how the relationship changes across elevations. P-values in bold are <0.05 to enhance visualisation of significant main effect of the trait and the interaction with elevation.

| **(a)** | | | | | |  | **(b)** | | | | | |  | **(c)** | | | | | |
| --- | --- | --- | --- | --- | --- | --- | --- | --- | --- | --- | --- | --- | --- | --- | --- | --- | --- | --- | --- |
| **Trait** | **Parameter** | **SS** | **Df** | **F-value** | **P-value** |  | **Trait** | **Parameter** | **SS** | **Df** | **F-value** | **P-value** |  | **Trait** | **Parameter** | **SS** | **Df** | **F-value** | **P-value** |
| Area | Intercept | 327.61 | 1 | 2382.22 | <0.001 |  | Area | Intercept | 35.6 | 1 | 127.98 | <0.001 |  | Area | Intercept | 0.16 | 1 | 14.23 | 0.0004 |
|  | **Plasticity** | 1.28 | 1 | 9.3 | **0.004** |  |  | Area | 0.1 | 1 | 0.06 | 0.808 |  |  | Area | 0 | 1 | 0.4 | 0.532 |
|  | Elevation | 4.91 | 2 | 17.84 | <0.001 |  |  | Elevation | 1.7 | 3 | 4.04 | 0.011 |  |  | Elevation | 0.09 | 3 | 2.57 | 0.062 |
|  | Plasticity × Elev. | 0.6 | 2 | 2.19 | 0.122 |  |  | Area × Elev. | 1.2 | 3 | 2.61 | 0.059 |  |  | Area × Elev. | 0.05 | 3 | 1.36 | 0.263 |
|  | Residuals | 7.01 | 51 |  |  |  |  | Residuals | 17.2 | 64 |  |  |  |  | Residuals | 0.73 | 64 |  |  |
| Nind | Intercept | 618.59 | 1 | 4566.16 | <0.001 |  | Nind | Intercept | 39.9 | 1 | 181.91 | <0.001 |  | Nind | Intercept | 0.22 | 1 | 25.52 | <0.001 |
|  | Plasticity | 0.29 | 1 | 2.14 | 0.15 |  |  | Indentation | 0 | 1 | 0.33 | 0.565 |  |  | Indentation | 0.01 | 1 | 1.21 | 0.275 |
|  | Elevation | 16.11 | 2 | 59.44 | <0.001 |  |  | Elevation | 8 | 3 | 13.1 | <0.001 |  |  | Elevation | 0.35 | 3 | 13.38 | <0.001 |
|  | **Plasticity × Elev.** | 0.97 | 2 | 3.59 | **0.035** |  |  | **Indentation × Elev.** | 2 | 3 | 3.24 | **0.028** |  |  | **Indentation × Elev.** | 0.2 | 3 | 7.64 | **<0.001** |
|  | Residuals | 6.91 | 51 |  |  |  |  | Residuals | 16.6 | 64 |  |  |  |  | Residuals | 0.56 | 64 |  |  |
| SLA | Intercept | 94.98 | 1 | 600.49 | <0.001 |  | SLA | Intercept | 74 | 1 | 335.44 | <0.001 |  | SLA | Intercept | 0.02 | 1 | 1.63 | 0.207 |
|  | Plasticity | 0.08 | 1 | 0.53 | 0.47 |  |  | **SLA** | 2.2 | 1 | 10.64 | **0.002** |  |  | **SLA** | 0.1 | 1 | 9.81 | **0.003** |
|  | Elevation | 5.9 | 2 | 18.65 | <0.001 |  |  | Elevation | 2.7 | 3 | 3.94 | 0.012 |  |  | Elevation | 0.01 | 3 | 0.2 | 0.893 |
|  | Plasticity × Elev. | 0.29 | 2 | 0.9 | 0.411 |  |  | SLA × Elev. | 1 | 3 | 1.15 | 0.338 |  |  | SLA × Elev. | 0.05 | 3 | 1.85 | 0.148 |
|  | Residuals | 8.07 | 51 |  |  |  |  | Residuals | 15.4 | 64 |  |  |  |  | Residuals | 0.63 | 64 |  |  |
| Chl | Intercept | 173.71 | 1 | 1152.03 | <0.001 |  | Chl | Intercept | 25.1 | 1 | 121.01 | <0.001 |  | Chl | Intercept | 0.27 | 1 | 32.66 | <0.001 |
|  | Plasticity | 0.53 | 1 | 3.49 | 0.067 |  |  | Chlorophyll | 0.2 | 1 | 0.68 | 0.414 |  |  | Chlorophyll | 0.04 | 1 | 4.42 | 0.039 |
|  | Elevation | 5.4 | 2 | 17.89 | <0.001 |  |  | Elevation | 9.5 | 3 | 14.05 | <0.001 |  |  | Elevation | 0.33 | 3 | 13.57 | <0.001 |
|  | Plasticity × Elev. | 0.05 | 2 | 0.15 | 0.86 |  |  | **Chlorophyll × Elev.** | 5 | 3 | 7.03 | **<0.001** |  |  | **Chlorophyll × Elev.** | 0.26 | 3 | 10.77 | **<0.001** |
|  | Residuals | 7.69 | 51 |  |  |  |  | Residuals | 12.7 | 64 |  |  |  |  | Residuals | 0.52 | 64 |  |  |
| Flav | Intercept | 294.05 | 1 | 1830.06 | <0.001 |  | Flav | Intercept | 23.4 | 1 | 87.03 | <0.001 |  | Flav | Intercept | 0.19 | 1 | 18.51 | <0.001 |
|  | Plasticity | 0.04 | 1 | 0.27 | 0.607 |  |  | Flavonol | 0.1 | 1 | 0.03 | 0.854 |  |  | Flavonol | 0.01 | 1 | 1 | 0.321 |
|  | Elevation | 24.59 | 2 | 76.52 | <0.001 |  |  | Elevation | 3.8 | 3 | 6.02 | 0.001 |  |  | Elevation | 0.17 | 3 | 5.43 | 0.002 |
|  | Plasticity × Elev. | 0.13 | 2 | 0.42 | 0.661 |  |  | Flavonol × Elev. | 1.1 | 3 | 2.09 | 0.110 |  |  | **Flavonol × Elev.** | 0.12 | 3 | 4.04 | **0.011** |
|  | Residuals | 8.19 | 51 |  |  |  |  | Residuals | 17.4 | 64 |  |  |  |  | Residuals | 0.66 | 64 |  |  |

**Table S5** Summary tables for significance tests of the regression slopes for the test of traits and their plasticity against fitness across elevations. Comparisons include: **(a)** trait values against fitness in **Fig. 5a**, **(b)** plasticity across elevations against fitness in **Fig. 5b**, **(c)** plasticity within elevations against fitness in **Fig. 6a**, and **(d)** plasticity within elevations against fitness in **Fig. 6b**. P-values in bold are <0.05 to aid visualisation. SE = 1 standard error, T = T-value relative to zero.

|  |  | **(a)** |  |  |  |  | **(b)** |  |  |  |  | **(c)** |  |  |  |  | **(d)** |  |  |  |
| --- | --- | --- | --- | --- | --- | --- | --- | --- | --- | --- | --- | --- | --- | --- | --- | --- | --- | --- | --- | --- |
| **Trait** | **Elevation** | **Slope** | **SE** | **T** | **P** |  | **Slope** | **SE** | **T** | **P** |  | **Slope** | **SE** | **T** | **P** |  | **Slope** | **SE** | **T** | **P** |
| Area | 500m | 1.03 | 0.2 | 4.90 | **<0.001** |  |  |  |  |  |  | 2.97 | 1.4 | 2.06 | **0.043** |  | -0.03 | 0.3 | -0.08 | 0.934 |
|  | 1000m | 0.68 | 0.2 | 3.92 | **<0.001** |  | 0.18 | 0.4 | 0.49 | 0.630 |  | -2.73 | 1.9 | -1.41 | 0.164 |  | 0.11 | 0.4 | 0.25 | 0.803 |
|  | 1500m | -0.15 | 0.4 | -0.42 | 0.676 |  | 0.55 | 0.3 | 1.66 | 0.103 |  | -1.17 | 1.3 | -0.93 | 0.354 |  | 0.23 | 0.3 | 0.82 | 0.415 |
|  | 2000m | -0.77 | 0.5 | -1.55 | 0.126 |  | 1.67 | 0.6 | 2.72 | **0.009** |  | 1.74 | 1.9 | 0.92 | 0.361 |  | -0.78 | 0.4 | -1.84 | 0.071 |
| Nind | 500m | -1.76 | 0.8 | -2.25 | **0.027** |  |  |  |  |  |  | -5.24 | 4.5 | -1.17 | 0.248 |  | 0.57 | 0.9 | 0.64 | 0.523 |
|  | 1000m | -0.93 | 1.0 | -0.90 | 0.370 |  | 1.34 | 1.1 | 1.26 | 0.213 |  | -2.87 | 2.6 | -1.09 | 0.280 |  | 0.46 | 0.5 | 0.89 | 0.376 |
|  | 1500m | -0.41 | 0.7 | -0.56 | 0.575 |  | -1.74 | 1.1 | -1.64 | 0.108 |  | -3.19 | 3.4 | -0.94 | 0.349 |  | 0.32 | 0.7 | 0.48 | 0.632 |
|  | 2000m | -0.97 | 0.7 | -1.39 | 0.170 |  | -2.06 | 0.8 | -2.66 | **0.010** |  | 7.33 | 2.9 | 2.51 | **0.015** |  | -2.84 | 0.6 | -4.93 | **<0.001** |
| SLA | 500m | -1.17 | 0.7 | -1.79 | 0.079 |  |  |  |  |  |  | -1.38 | 2.3 | -0.61 | 0.544 |  | 0.41 | 0.5 | 0.83 | 0.408 |
|  | 1000m | -1.14 | 0.6 | -2.01 | **0.049** |  | -1.65 | 1.2 | -1.37 | 0.176 |  | -6.05 | 2.7 | -2.26 | **0.027** |  | 0.52 | 0.6 | 0.89 | 0.375 |
|  | 1500m | 1.54 | 0.9 | 1.73 | 0.088 |  | 0.67 | 1.2 | 0.54 | 0.590 |  | -3.54 | 3.4 | -1.03 | 0.309 |  | 0.79 | 0.7 | 1.06 | 0.295 |
|  | 2000m | 3.33 | 1.5 | 2.28 | **0.026** |  | -0.60 | 1.3 | -0.46 | 0.647 |  | -7.46 | 2.8 | -2.67 | **0.010** |  | 2.11 | 0.6 | 3.5 | **0.001** |
| Chl | 500m | 1.59 | 0.6 | 2.53 | **0.014** |  |  |  |  |  |  | -9.88 | 2.9 | -3.46 | **0.001** |  | 0.25 | 0.6 | 0.42 | 0.676 |
|  | 1000m | 0.92 | 0.6 | 1.68 | 0.098 |  | 1.32 | 0.9 | 1.44 | 0.155 |  | -2.72 | 2.3 | -1.2 | 0.235 |  | 0.39 | 0.5 | 0.82 | 0.416 |
|  | 1500m | 0.75 | 0.8 | 1.00 | 0.321 |  | 1.51 | 0.9 | 1.64 | 0.107 |  | -2.03 | 1.9 | -1.05 | 0.299 |  | 0.74 | 0.4 | 1.83 | 0.071 |
|  | 2000m | -0.66 | 0.8 | -0.78 | 0.437 |  | 0.63 | 1.3 | 0.48 | 0.634 |  | 10.25 | 3.4 | 3.06 | **0.003** |  | -3.69 | 0.7 | -5.3 | **<0.001** |
| Flav | 500m | 1.63 | 0.6 | 2.92 | **0.005** |  |  |  |  |  |  | -4.56 | 3.7 | -1.22 | 0.226 |  | -0.04 | 0.8 | -0.06 | 0.955 |
|  | 1000m | 1.10 | 0.5 | 2.09 | **0.041** |  | 0.18 | 1.0 | 0.18 | 0.861 |  | -1.57 | 2.7 | -0.58 | 0.563 |  | 0.19 | 0.6 | 0.34 | 0.737 |
|  | 1500m | -0.65 | 0.6 | -1.17 | 0.246 |  | -0.12 | 0.7 | -0.17 | 0.867 |  | -2.28 | 2.9 | -0.79 | 0.435 |  | 0.88 | 0.6 | 1.45 | 0.152 |
|  | 2000m | -1.41 | 0.6 | -2.45 | **0.017** |  | -0.79 | 0.6 | -1.22 | 0.227 |  | 7.20 | 3.6 | 2.00 | **0.050** |  | -2.39 | 0.8 | -3.18 | **0.002** |

**Table S6** Summary tables for pairwise significance tests of the regression slopes between elevation for the test of traits and their plasticity against fitness. Comparisons include: **(a)** trait values against fitness in **Fig. 5a**, **(b)** plasticity across elevations against fitness in **Fig. 5b**, **(c)** plasticity within elevations against fitness in **Fig. 6a**, and **(d)** plasticity within elevations against fitness in **Fig. 6b**. P-values in bold are <0.05 to aid visualisation. SE = 1 standard error, T = T-value relative to zero.

|  |  | **(a)** |  |  |  |  | **(b)** |  |  |  |  | **(c)** |  |  |  |  | **(d)** |  |  |  |
| --- | --- | --- | --- | --- | --- | --- | --- | --- | --- | --- | --- | --- | --- | --- | --- | --- | --- | --- | --- | --- |
| **Trait** | **Pairwise test** | **Diff.** | **SE** | **T** | **P** |  | **Diff.** | **SE** | **T** | **P** |  | **Diff.** | **SE** | **T** | **P** |  | **Diff.** | **SE** | **T** | **P** |
| Area | 500m-1000m | 0.35 | 0.3 | 1.29 | 0.575 |  |  |  |  |  |  | 5.70 | 2.4 | 2.36 | 0.095 |  | -0.14 | 0.5 | -0.25 | 0.994 |
|  | 500m-1500m | 1.18 | 0.4 | 2.83 | **0.031** |  |  |  |  |  |  | 4.14 | 1.9 | 2.17 | 0.144 |  | -0.26 | 0.4 | -0.60 | 0.931 |
|  | 500m-2000m | 1.80 | 0.5 | 3.32 | **0.008** |  |  |  |  |  |  | 1.23 | 2.4 | 0.52 | 0.955 |  | 0.75 | 0.5 | 1.41 | 0.497 |
|  | 1000m-1500m | 0.83 | 0.4 | 2.07 | 0.172 |  | -0.38 | 0.5 | -0.77 | 0.726 |  | -1.56 | 2.3 | -0.68 | 0.906 |  | -0.12 | 0.5 | -0.24 | 0.995 |
|  | 1000m-2000m | 1.45 | 0.5 | 2.74 | **0.038** |  | -1.49 | 0.7 | -2.09 | 0.101 |  | -4.48 | 2.7 | -1.65 | 0.358 |  | 0.89 | 0.6 | 1.46 | 0.465 |
|  | 1500m-2000m | 0.62 | 0.6 | 1.01 | 0.744 |  | -1.11 | 0.7 | -1.60 | 0.256 |  | -2.92 | 2.3 | -1.28 | 0.578 |  | 1.01 | 0.5 | 1.98 | 0.205 |
| Nind | 500m-1000m | -0.84 | 1.3 | -0.65 | 0.915 |  |  |  |  |  |  | -2.36 | 5.2 | -0.45 | 0.969 |  | 0.11 | 1.0 | 0.10 | 1.000 |
|  | 500m-1500m | -1.35 | 1.1 | -1.27 | 0.588 |  |  |  |  |  |  | -2.04 | 5.6 | -0.36 | 0.983 |  | 0.25 | 1.1 | 0.22 | 0.996 |
|  | 500m-2000m | -0.79 | 1.1 | -0.75 | 0.875 |  |  |  |  |  |  | -12.57 | 5.4 | -2.35 | 0.098 |  | 3.41 | 1.1 | 3.23 | **0.010** |
|  | 1000m-1500m | -0.52 | 1.3 | -0.41 | 0.977 |  | 3.08 | 1.5 | 2.05 | 0.111 |  | 0.32 | 4.3 | 0.07 | 1.000 |  | 0.14 | 0.8 | 0.17 | 0.998 |
|  | 1000m-2000m | 0.05 | 1.2 | 0.04 | 1.000 |  | 3.40 | 1.3 | 2.59 | **0.033** |  | -10.20 | 3.9 | -2.59 | 0.056 |  | 3.30 | 0.8 | 4.26 | **<0.001** |
|  | 1500m-2000m | 0.56 | 1.0 | 0.55 | 0.945 |  | 0.32 | 1.3 | 0.24 | 0.968 |  | -10.52 | 4.5 | -2.36 | 0.096 |  | 3.16 | 0.9 | 3.59 | **0.004** |
| SLA | 500m-1000m | -0.03 | 0.9 | -0.03 | 1.000 |  |  |  |  |  |  | 4.67 | 3.5 | 1.34 | 0.544 |  | -0.11 | 0.8 | -0.15 | 0.999 |
|  | 500m-1500m | -2.71 | 1.1 | -2.45 | 0.077 |  |  |  |  |  |  | 2.16 | 4.1 | 0.52 | 0.953 |  | -0.38 | 0.9 | -0.43 | 0.974 |
|  | 500m-2000m | -4.50 | 1.6 | -2.81 | **0.032** |  |  |  |  |  |  | 6.08 | 3.6 | 1.69 | 0.335 |  | -1.71 | 0.8 | -2.20 | 0.134 |
|  | 1000m-1500m | -2.68 | 1.1 | -2.54 | 0.063 |  | -2.32 | 1.7 | -1.34 | 0.378 |  | -2.51 | 4.4 | -0.58 | 0.939 |  | -0.27 | 0.9 | -0.29 | 0.992 |
|  | 1000m-2000m | -4.47 | 1.6 | -2.85 | **0.029** |  | -1.05 | 1.8 | -0.59 | 0.824 |  | 1.41 | 3.9 | 0.36 | 0.983 |  | -1.60 | 0.8 | -1.91 | 0.233 |
|  | 1500m-2000m | -1.78 | 1.7 | -1.04 | 0.724 |  | 1.27 | 1.8 | 0.71 | 0.760 |  | 3.92 | 4.4 | 0.88 | 0.813 |  | -1.33 | 1.0 | -1.39 | 0.513 |
| Chl | 500m-1000m | 0.67 | 0.8 | 0.80 | 0.855 |  |  |  |  |  |  | -7.16 | 3.6 | -1.96 | 0.212 |  | -0.14 | 0.8 | -0.18 | 0.998 |
|  | 500m-1500m | 0.84 | 1.0 | 0.86 | 0.827 |  |  |  |  |  |  | -7.85 | 3.4 | -2.28 | 0.114 |  | -0.49 | 0.7 | -0.68 | 0.904 |
|  | 500m-2000m | 2.25 | 1.1 | 2.14 | 0.151 |  |  |  |  |  |  | -20.13 | 4.4 | -4.57 | **<0.001** |  | 3.94 | 0.9 | 4.31 | **<0.001** |
|  | 1000m-1500m | 0.17 | 0.9 | 0.19 | 0.998 |  | -0.19 | 1.3 | -0.15 | 0.988 |  | -0.69 | 3.0 | -0.23 | 0.996 |  | -0.35 | 0.6 | -0.57 | 0.942 |
|  | 1000m-2000m | 1.59 | 1.0 | 1.57 | 0.401 |  | 0.68 | 1.6 | 0.43 | 0.905 |  | -12.97 | 4.0 | -3.21 | **0.011** |  | 4.08 | 0.8 | 4.85 | **<0.001** |
|  | 1500m-2000m | 1.41 | 1.1 | 1.25 | 0.598 |  | 0.88 | 1.6 | 0.54 | 0.850 |  | -12.28 | 3.9 | -3.17 | **0.012** |  | 4.43 | 0.8 | 5.51 | **<0.001** |
| Flav | 500m-1000m | 0.52 | 0.8 | 0.68 | 0.904 |  |  |  |  |  |  | -2.99 | 4.6 | -0.65 | 0.915 |  | -0.23 | 1.0 | -0.24 | 0.995 |
|  | 500m-1500m | 2.27 | 0.8 | 2.90 | **0.026** |  |  |  |  |  |  | -2.28 | 4.7 | -0.48 | 0.963 |  | -0.93 | 1.0 | -0.94 | 0.785 |
|  | 500m-2000m | 3.04 | 0.8 | 3.79 | **0.002** |  |  |  |  |  |  | -11.77 | 5.2 | -2.27 | 0.117 |  | 2.35 | 1.1 | 2.16 | 0.144 |
|  | 1000m-1500m | 1.75 | 0.8 | 2.29 | 0.111 |  | 0.30 | 1.3 | 0.24 | 0.969 |  | 0.72 | 4.0 | 0.18 | 0.998 |  | -0.69 | 0.8 | -0.83 | 0.838 |
|  | 1000m-2000m | 2.52 | 0.8 | 3.21 | **0.011** |  | 0.97 | 1.2 | 0.80 | 0.706 |  | -8.77 | 4.5 | -1.95 | 0.218 |  | 2.58 | 0.9 | 2.74 | **0.038** |
|  | 1500m-2000m | 0.77 | 0.8 | 0.96 | 0.772 |  | 0.67 | 1.0 | 0.70 | 0.764 |  | -9.49 | 4.6 | -2.05 | 0.181 |  | 3.27 | 1.0 | 3.38 | **0.007** |
